## Supplementary material for "Partial observation can induce mechanistic mismatches in data-constrained models of neural dynamics": Code: bptt.py

```
import numpy as np
import torch
import torch.nn as nn
import torch.nn.functional as F
from torch.optim import SGD
from torch.utils.data import Dataset, DataLoader
import matplotlib.pyplot as plt
import pickle as pk
import os
os.environ["KMP_DUPLICATE_LIB_OK"]="TRUE"

conditions_path = './saved_conditions'
activity_path = './saved_activity'

class MyDataset(Dataset):
    def __init__(self, data, window_size):
        self.data = data
        self.window_size = window_size

    def __getitem__(self, index):
        macro_idx = index // 2
        offset = 0 if index % 2 else self.window_size // 2
        x = self.data[:, macro_idx * self.window_size + offset: (macro_idx + 1) * self.window_size + offset]
        return x

    def __len__(self):
        return 2 * self.data.shape[1] // self.window_size - 2
class RNN(nn.Module):
    def __init__(self, N_neurons, N_obs, alpha = 0.01, beta=0.125, f = nn.Identity, A = None, use_stochastic_tf = True):
        super(RNN, self).__init__()
        if A is None:
            self.w = nn.Parameter(torch.randn(N_neurons, N_neurons) * (0.1/ np.sqrt(N_neurons)), requires_grad=True)
        else:
            self.w = A
        self.alpha = alpha
        self.beta = beta
        self.N_neurons = N_neurons
        self.N_obs = N_obs
        self.f = f()
        self.use_stochastic_tf = use_stochastic_tf
        self.extra_ic = torch.zeros((self.N_neurons - self.N_obs))
    def forward_model(self, x):
        return x + self.alpha * (-x + ((self.w @ self.f(x).T).T))

    def forward(self, x_teacher, T_steps):
        bs = x_teacher.shape[0]
        out = []
        if len(x_teacher.shape) == 2:
            ic = x_teacher
            x_teacher = None
        else:
            ic = x_teacher[:, :, 0] #bs, n_obs
        extra_ic = self.extra_ic.unsqueeze(0)
        x = torch.cat([ic, extra_ic.expand(bs, extra_ic.shape[1])], dim = 1) # init, bs x N_neurons
        out.append(x)
        x = self.forward_model(x)
        for t in range(1, T_steps):
            x_gtf = self.generalized_teacher_forcing(x, x_teacher[:,:, t] if x_teacher is not None else None)
            x = self.forward_model(x_gtf)
            out.append(x)
        out = torch.stack(out) # nt, bs, N_neurons
        return out.permute(1,2,0) #bs, N_neurons, nt
    def generalized_teacher_forcing(self, x, x_tch):
        if x_tch is not None:
            x_tch = torch.cat([x_tch, x[:, x_tch.shape[1]:]], dim=-1)
        else:
            return x
        if self.use_stochastic_tf:
            if np.random.rand() < self.beta:
                return x_tch
            else:
                return x
        else:
            return self.beta * x_tch + (1-self.beta) * x
def loss_function(model, x_obs):
    x_obs_hat = model(x_obs, x_obs.shape[-1])[:, :x_obs.shape[1], :]
    return F.mse_loss(model.f(x_obs_hat), model.f(x_obs), reduction='sum')

def train_RNN(gt_activity, N_obs=50, N_hidden=0, epochs=200, T=1000, T_tot = 1000, lr=1e-3, beta = 0.0, f_suffix=''):
    '''

    :param gt_activity: data to fit
    :param N_obs: observed neurons to fit
    :param N_hidden: hidden neurons to add
    :param epochs:
    :param T: length of bptt truncation
    :param T_tot: total length of activity to fit
    :param lr: learning rate
    :param beta: teacher forcing ratio
    :return:
    '''
    rnn_student = RNN(N_obs + N_hidden, N_obs, alpha = 0.01, beta=beta, f = nn.Tanh)
    dataset = MyDataset(gt_activity[0], window_size=T)
    loader = DataLoader(dataset, batch_size=16, shuffle=True)
    opt = SGD(rnn_student.parameters(), lr=lr, momentum=0.9)
    batch_idx = 0
    for i in range(epochs):
        for batch in loader:
            batch_idx += 1
            opt.zero_grad()
            loss = loss_function(rnn_student, batch[:, :N_obs])
            loss.backward()
            opt.step()
            print('epoch:', i, 'batch:', batch_idx,'loss:', loss)
    student_w = rnn_student.w.detach().numpy()
    bptt_weights_f = open(os.path.join(conditions_path, 'bptt_weights_{}.npy'.format(f_suffix)), 'wb')
    np.save(bptt_weights_f, student_w)

if __name__ == "__main__":
    load_from_file = True
    if len(os.listdir(conditions_path)) == 0:
        print('No condition files found.')
        load_from_file = False
    if load_from_file:
        gt_activity_from_file = np.load(os.path.join(activity_path, 'triangular_50_teacher_activity.npy'))
        gt_activity = torch.from_numpy(gt_activity_from_file).T.unsqueeze(0).float()
        train_RNN(gt_activity, 50,  N_hidden=0, T=500, T_tot=30000, epochs=200, lr=0.01, beta = 0.5)
```
