## Supplementary material for "Partial observation can induce mechanistic mismatches in data-constrained models of neural dynamics": Code: hidden_neuron_plots.py

```
import os
import pickle as pk

import numpy as np
from matplotlib import pyplot as plt


from make_RNN_plots import create_fit_plots
from simulation_conditions import generate_simulation_conditions

# Make a folder to store figures
fig_path = './figures'
if not os.path.isdir(fig_path):
    os.makedirs(fig_path)
conditions_path = './saved_conditions'
activity_path = './saved_activity'
print(f"Storing figures in {fig_path}")

if __name__ == "__main__":
    load_from_file = True
    if len(os.listdir(conditions_path)) == 0:
        print('No condition files found.')
        load_from_file = False

    '''Generic non-normal teacher, nonlinear, MAP'''
    f_name = 'triangular'
    if load_from_file:
        conditions = pk.load(open(os.path.join(conditions_path, f_name + '.pk'), 'rb'))
    else:
        conditions = generate_simulation_conditions(500, 30000, 25, network='triangular', ortho_transform = True, diagonal_comp=0.0, use_schur=True, scale = 2.8)

    n_obs = 50
    nt = 30000
    conditions_map = conditions.copy()
    conditions_map['obs_noise'] = np.hstack(
        [conditions['obs_noise'][:, :n_obs], np.zeros((nt, max(0, n_obs - conditions['obs_noise'].shape[1])))])
    f_name = f_name + '_{}'.format(n_obs)

    create_fit_plots(500, nt, n_obs, v1=0.2, rho=0.001, fit_method='direct', conditions=conditions_map,
                     phi=lambda x: np.tanh(x), use_direct_projection=False, f_suffix=f_name,
                     nonlinear_weights=False, plot_time=16000)

    '''BPTT'''
    f_name = 'triangular'
    n_obs = 50
    nt = 30000
    conditions_bptt = conditions.copy()
    conditions_bptt['obs_noise'] = np.hstack([conditions['obs_noise'][:, :n_obs], np.zeros((nt, max(0, n_obs - conditions['obs_noise'].shape[1])))])
    f_name = f_name+'_{}'.format(n_obs)
    if load_from_file:
        file = 'bptt_weights_50_hidden_triangular.npy'
        create_fit_plots(500, nt, n_obs, v1=0.2, rho=0.001, fit_method='file', conditions=conditions_bptt,
                         phi=lambda x: np.tanh(x), use_direct_projection=False, f_suffix=f_name + '_bptt',
                         nonlinear_weights=False, plot_time=16000, load_from_file=file)

    '''BPTT + hidden'''
    n_obs = 250
    nt = 30000
    conditions_bptt = conditions.copy()
    conditions_bptt['obs_noise'] = np.hstack([conditions['obs_noise'][:, :n_obs], np.zeros((nt, max(0, n_obs - conditions['obs_noise'].shape[1])))])
    f_name = f_name+'_{}'.format(n_obs)
    if load_from_file:
        file = 'bptt_weights_50_200_hidden_triangular.npy'
        create_fit_plots(500, nt, n_obs, v1=0.2, rho=0.001, fit_method='file', conditions=conditions_bptt,
                         phi=lambda x: np.tanh(x), use_direct_projection=False, f_suffix=f_name + '_bptt',
                         nonlinear_weights=False, plot_time=16000, load_from_file=file)

    plt.show()
```
