## Supplementary material for "Partial observation can induce mechanistic mismatches in data-constrained models of neural dynamics": Code: integration_plots.py

```
import os
import pickle as pk

import numpy as np
from matplotlib import pyplot as plt

from RNN import RNN
from RNN_fit import fit_rslds
from make_RNN_plots import make_input_readout_plots, plot_rslds_fit_results
from simulation_conditions import generate_simulation_conditions

# Make a folder to store figures
fig_path = './figures'
if not os.path.isdir(fig_path):
    os.makedirs(fig_path)
conditions_path = './saved_conditions'
activity_path = './saved_activity'
print(f"Storing figures in {fig_path}")

if __name__ == "__main__":
    load_from_file = True
    if len(os.listdir(conditions_path)) == 0:
        print('No condition files found.')
        load_from_file = False

    ''' line attractor integration '''
    f_name_la = 'la_integrator'
    if load_from_file:
        conditions = pk.load(open(os.path.join(conditions_path, f_name_la+'.pk'), 'rb'))
    else:
        conditions = generate_simulation_conditions(500, 5000, 25, network='lr_n', rank=500, max_eval=0.999, min_eval = 0.2, diagonal=True, similarity_transform = True)

    rnn = RNN(500, 5000, 0.01, 0.1, 0.2, 25, 0, conditions=conditions, integrate_sum=False)
    make_input_readout_plots(rnn, f_name_la, subplot=True)

    if load_from_file:
        rslds, q_elbos_lem, zhat_lem, xhat_lem, yhat_lem = pk.load(open(os.path.join(conditions_path, f_name_la + '_rslds.pk'), 'rb'))
    else:
        input = rnn.inp.copy()
        input /= np.max(np.abs(input))
        rslds, q_elbos_lem, zhat_lem, xhat_lem, yhat_lem = fit_rslds(rnn.y, 25, 1, 5, n_iters=200, input=input)

    plot_rslds_fit_results(rnn, rslds, q_elbos_lem, zhat_lem, xhat_lem, yhat_lem, f_suffix=f_name_la,
                           activity_subplots=True)

    ''' functionally feedforward integration '''
    f_name_nn = 'nn_integrator'
    if load_from_file:
        conditions = pk.load(open(os.path.join(conditions_path, f_name_nn + '.pk'), 'rb'))
    else:
        conditions = generate_simulation_conditions(500, 5000, 25, network = 'delay', skip_scale = 0.5, skip_interval = 1, ortho_transform = True)

    rnn = RNN(500, 5000, 0.01, 0.1, 0.2, 25, 0, conditions=conditions, integrate_sum=True)
    make_input_readout_plots(rnn, f_name_la, subplot=True)

    if load_from_file:
        rslds, q_elbos_lem, zhat_lem, xhat_lem, yhat_lem = pk.load(
            open(os.path.join(conditions_path, f_name_nn + '_rslds.pk'), 'rb'))
    else:
        input = rnn.inp.copy()
        input /= np.max(np.abs(input))
        rslds, q_elbos_lem, zhat_lem, xhat_lem, yhat_lem = fit_rslds(rnn.y, 25, 1, 5, n_iters=200, input=input)

    plot_rslds_fit_results(rnn, rslds, q_elbos_lem, zhat_lem, xhat_lem, yhat_lem, f_suffix=f_name_nn,
                           activity_subplots=True, ff_chain=True)
    plt.show()
```
