## Supplementary material for "Partial observation can induce mechanistic mismatches in data-constrained models of neural dynamics": Code: low_rank_plots.py

```
import os
import pickle as pk

import numpy as np
from matplotlib import pyplot as plt


from make_RNN_plots import create_fit_plots
from simulation_conditions import generate_simulation_conditions

# Make a folder to store figures
fig_path = './figures'
if not os.path.isdir(fig_path):
    os.makedirs(fig_path)
conditions_path = './saved_conditions'
activity_path = './saved_activity'
print(f"Storing figures in {fig_path}")

if __name__ == "__main__":
    load_from_file = True
    if len(os.listdir(conditions_path)) == 0:
        print('No condition files found.')
        load_from_file = False

    '''Rank 2 linear teacher'''
    f_name = 'lr_nn_rank_2_linear'
    if load_from_file:
        conditions = pk.load(open(os.path.join(conditions_path, f_name + '.pk'), 'rb'))
    else:
        conditions = generate_simulation_conditions(500, 30000, 25, network = 'lr_nn', rank = 2, scale = 0.2)
    create_fit_plots(500, 30000, 25, v1=0.2, rho=0.001, fit_method='direct', conditions=conditions,
                     phi=lambda x: x, use_direct_projection=False, f_suffix=f_name,
                     nonlinear_weights=False, plot_time=4000, data=None)

    '''Rank 3 linear teacher'''
    f_name = 'lr_nn_rank_3_linear'
    if load_from_file:
        conditions = pk.load(open(os.path.join(conditions_path, f_name + '.pk'), 'rb'))
    else:
        conditions = generate_simulation_conditions(500, 30000, 25, network = 'lr_nn', rank = 3, scale = 0.2)
    create_fit_plots(500, 30000, 25, v1=0.2, rho=0.001, fit_method='direct', conditions=conditions,
                     phi=lambda x: x, use_direct_projection=False, f_suffix=f_name,
                     nonlinear_weights=False, plot_time=4000, data=None)

    '''Rank 2 nonlinear teacher with oscillatory dynamics'''
    f_name = 'lr_nn_rank_2_nonlinear_osc'
    if load_from_file:
        conditions = pk.load(open(os.path.join(conditions_path, f_name + '.pk'), 'rb'))
    else:
        # depending on instantiation, conditions below could support learning spurious FPs, LCs, or nothing unusual
        conditions = generate_simulation_conditions(500, 30000, 25, network='lr_nn_osc', shift_scale=12)

    # pad saved conditions to support simulating larger students
    n_obs = 50
    nt = 30000
    conditions_map = conditions.copy()
    conditions_map['obs_noise'] = np.hstack([conditions['obs_noise'][:, :n_obs], np.zeros((nt, max(0, n_obs - conditions['obs_noise'].shape[1])))])
    f_name = f_name+'_{}'.format(n_obs)

    create_fit_plots(500, nt, n_obs, v1=0.2, rho=0.001, fit_method='direct', conditions=conditions_map,
                     phi=lambda x: np.tanh(x), use_direct_projection=False, f_suffix=f_name,
                     nonlinear_weights=False, plot_time=8000)

    '''CORNN'''
    f_name = 'lr_nn_rank_2_nonlinear_osc'
    n_obs = 25
    f_name = f_name + '_{}'.format(n_obs)
    create_fit_plots(500, nt, n_obs, v1=0.2, rho=0.001, fit_method='cornn', conditions=conditions,
                     phi=lambda x: np.tanh(x), use_direct_projection=False, f_suffix=f_name+'_cornn',
                     nonlinear_weights=True, plot_time=8000)

    '''FORCE'''
    f_name = 'lr_nn_rank_2_nonlinear_osc'

    n_obs = 50
    nt = 30000
    conditions_force = conditions.copy()
    conditions_force['obs_noise'] = np.hstack([conditions['obs_noise'][:, :n_obs], np.zeros((nt, max(0, n_obs - conditions['obs_noise'].shape[1])))])
    f_name = f_name+'_{}'.format(n_obs)
    if load_from_file:
        file = 'weights_lr_nn_rank_2_nonlinear_osc_{}_force.npy'.format(n_obs)
        create_fit_plots(500, nt, n_obs, v1=0.2, rho=0.001, fit_method='file', conditions=conditions_force,
                         phi=lambda x: np.tanh(x), use_direct_projection=False, f_suffix=f_name+'_force',
                         nonlinear_weights=False, plot_time=8000, load_from_file=file)
    else:
        create_fit_plots(500, nt, n_obs, v1=0.2, rho=0.001, fit_method='force', conditions=conditions_force,
                         phi=lambda x: np.tanh(x), use_direct_projection=False, f_suffix=f_name+'_force',
                         nonlinear_weights=False, plot_time=8000)

    '''BPTT'''
    f_name = 'lr_nn_rank_2_nonlinear_osc'
    n_obs = 50
    nt = 30000
    conditions_bptt = conditions.copy()
    conditions_bptt['obs_noise'] = np.hstack([conditions['obs_noise'][:, :n_obs], np.zeros((nt, max(0, n_obs - conditions['obs_noise'].shape[1])))])
    f_name = f_name+'_{}'.format(n_obs)

    if load_from_file:
        file = 'weights_lr_nn_rank_2_nonlinear_osc_{}_bptt.npy'.format(n_obs)
        create_fit_plots(500, nt, n_obs, v1=0.2, rho=0.001, fit_method='file', conditions=conditions_bptt,
                         phi=lambda x: np.tanh(x), use_direct_projection=False, f_suffix=f_name + '_bptt',
                         nonlinear_weights=False, plot_time=8000, load_from_file=file)
    plt.show()
```
