## Supplementary material for "Partial observation can induce mechanistic mismatches in data-constrained models of neural dynamics": Code: make_RNN_plots.py

```
from sklearn.decomposition import PCA
from RNN_fit import *
import os
from RNN import *
import matplotlib.pyplot as plt
from mpl_toolkits.mplot3d.art3d import Line3DCollection

# Make a folder to store figures
fig_path = './figures'
if not os.path.isdir(fig_path):
    os.makedirs(fig_path)
conditions_path = './saved_conditions'
activity_path = './saved_activity'
print(f"Storing figures in {fig_path}")

def make_single_neuron_traces(gt, learned, alpha=0.01, rescale=0.7, trace_sep=2.5, f_suffix=''):
    fig, ax = plt.subplots(1, 1, figsize=(3, 3), dpi=150)
    T, N = gt.shape
    neuron_plot_idxes = [4,5,6,7,8]
    t = np.arange(T) * alpha
    for idx, i in enumerate(neuron_plot_idxes):
        #ax.axhline(y=trace_sep * idx, color = 'gray', linestyle='--', linewidth=0.5)
        ax.plot(t, rescale * gt[:, i] + trace_sep * idx, color='r')
        ax.plot(t, rescale * learned[:, i] + trace_sep * idx, color='b')
    ax.set_yticks([])
    ax.set_ylabel('Neural Activity (A.U.)', fontsize=15)
    ax.set_xlabel('t', fontsize=15)
    ax.spines[['right', 'top']].set_visible(False)
    ax.set_xticks([0, int(T * alpha)])
    ax.xaxis.set_tick_params(labelsize=15)
    plt.tight_layout()
    plt.savefig(os.path.join(fig_path, 'neuron_traces_{}.pdf'.format(f_suffix)))

def create_fit_plots(n_neuron, nt, n_obs, alpha=0.01, v0=0, v1 = 0.2, s_obs = 0.0, conditions = None, network = 'gaussian', phi = lambda x: x, fit_method='direct', rho = 0.0,
                     use_direct_projection=False, nonlinear_weights = False, f_suffix='',plot_time=8000, plot_flowfields=False, load_from_file=None, **kwargs):
    if conditions is None:
        conditions = generate_simulation_conditions(n_neuron, nt, n_obs, network = network, nonlinear_weights=nonlinear_weights, **kwargs)
    else:
        conditions = conditions.copy()
    conditions['obs_noise'] = conditions['obs_noise'][:nt]
    conditions['inp_noise'] = conditions['inp_noise'][:nt]
    rnn = RNN(n_neuron, nt, alpha, v0, v1, n_obs, s_obs, phi = phi, conditions = conditions, nonlinear_weights=nonlinear_weights)
    rnn_activity_f = os.path.join(activity_path, f_suffix + '_teacher_activity.npy')
    np.save(rnn_activity_f,rnn.x)
    noise = conditions['inp_noise'].T * v1
    if fit_method == 'implicit':
        A = fit_solution_implicit(rnn.w, rnn.x.T, n_obs, alpha, rho, noise, phi=phi, noise_limit=False)
    elif fit_method == 'long_time_limit':
        A = fit_solution_implicit(rnn.w, rnn.x.T, n_obs, alpha, rho, noise, phi=phi, noise_limit=True)
    elif fit_method == 'cornn':
        A = fit_cornn(rnn.y, inputs=rnn.inp)
        A = A[:n_obs].T
    elif fit_method == 'direct':
        A = fit_solution_direct(rnn.y.T, n_obs, alpha, rho, phi=phi)
    elif fit_method == 'gd':
        A = fit_solution_grad_descent(rnn.y.T, phi=phi)
    elif fit_method == 'force':
        train_out = trainMultiRegionRNN(rnn.y.T, tauRNN=1, dtData=alpha, ampInWN=0.0, P0=0.01, nonLinearity=phi, nRunTrain=1000, g=1.5)
        A = train_out['J']
        # weights_f = os.path.join(conditions_path, 'force_weights_50_{}.npy'.format(f_suffix))
        # np.save(weights_f, A)
    elif fit_method == 'file':
        assert load_from_file is not None, "No file given!"
        assert os.path.isfile(os.path.join(conditions_path, load_from_file)), "no such file!"
        A = np.load(os.path.join(conditions_path, load_from_file))

    conditions_new = {'ic': conditions['ic'][:n_obs],
                      'w': A,
                      'inp_noise': conditions['inp_noise'][:, :n_obs],
                      'obs_noise': conditions['obs_noise'][:, :n_obs],
                      'U': conditions['U'][:n_obs] if conditions['U'] is not None else None}

    if fit_method in ['force', 'bptt']:
        v1 = 0
    rnn_student = RNN(n_obs, nt, alpha, v0, v1, n_obs, s_obs, phi=phi if fit_method != 'cornn' else lambda x: np.tanh(x),
                      conditions = conditions_new, nonlinear_weights=nonlinear_weights if fit_method != 'cornn' else True)
    if not plot_flowfields:
        make_single_neuron_traces(rnn.x[:plot_time, :n_obs], rnn_student.x[:plot_time], f_suffix=f_suffix)
        get_rnn_limit_sets(A,
                           use_direct_projection=use_direct_projection, transient_frac=0.05, phi = phi,
                           fname='limit_set_fit_{}.pdf'.format(f_suffix), nonlinear_weights=nonlinear_weights)

        get_rnn_limit_sets(rnn.w, use_direct_projection=use_direct_projection, transient_frac=0.06, phi = phi,
                           nonlinear_weights=nonlinear_weights, fname='limit_set_gt_{}.pdf'.format(f_suffix))
    else:
        make_pca_plot(rnn.x, transient_frac=0.0, use_3d=False, w=rnn.w, title='Ground Truth',
                      save_name='gt_flowfield_{}.pdf'.format(f_suffix), flow_scale=150, bbox_lims=True)
        make_pca_plot(rnn_student.x, transient_frac=0.0, use_3d=False, w=A, title='Learned',
                      save_name='fit_flowfield_{}.pdf'.format(f_suffix), flow_scale=50, bbox_lims = True)

    make_eigenvalue_plot({'Ground Truth': rnn.w, 'Learned': A}, f_suffix=f_suffix)

    return conditions

# set steps to 10000 for longer iteration
def get_rnn_limit_sets(w, steps=5000, use_direct_projection=False,
                       dt=0.01, transient_frac=0.01, n_traj = 15, plot_interval=4, phi = lambda x: x, nonlinear_weights = False, radius = 0.5, fname='limit_set.pdf'):
    N = len(w)
    # turn off noise for limit sets
    inp_noise = np.zeros((steps, N))
    obs_noise = np.zeros((steps, N))
    data_ranges = [[0, 0], [0, 0], [0, 0]]
    x_all = []
    x0_all = np.random.normal(0,1,(n_traj, N)) * radius
    limit_set_f = os.path.join(conditions_path, fname+'_limit_set_init.npy')
    if os.path.isfile(limit_set_f):
        with open(limit_set_f,'rb') as f:
            x0_all = np.load(f)
    else:
        with open(limit_set_f, 'wb') as f:
            np.save(f, x0_all)
    for i in range(n_traj):
        x0 = x0_all[i]
        conditions = {'ic':x0, 'w':w, 'inp_noise':inp_noise, 'obs_noise':obs_noise, 'U': None}
        rnn = RNN(N, steps, dt, 0,0,N, 0, phi=phi, conditions = conditions, nonlinear_weights=nonlinear_weights)
        x_all.append(rnn.x[::plot_interval])
    if N > 2:
        make_pca_plot(x_all, use_3d=True, use_direct_projection=use_direct_projection, transient_frac=transient_frac,
        data_ranges=data_ranges)
    plt.savefig(os.path.join(fig_path, fname))

def set_axes_equal_range(X_pca, ranges, ax: plt.Axes, scale=1):
    """Set 3D plot axes to equal scale.

    Make axes of 3D plot have equal scale so that spheres appear as
    spheres and cubes as cubes.  Required since `ax.axis('equal')`
    and `ax.set_aspect('equal')` don't work on 3D.
    """
    ranges[0][0] = min(ranges[0][0], X_pca[:, 0].min())
    ranges[0][1] = max(ranges[0][1], X_pca[:, 0].max())
    ranges[1][0] = min(ranges[1][0], X_pca[:, 1].min())
    ranges[1][1] = max(ranges[1][1], X_pca[:, 1].max())
    ranges[2][0] = min(ranges[2][0], X_pca[:, 2].min())
    ranges[2][1] = max(ranges[2][1], X_pca[:, 2].max())
    max_range = scale * np.array(
        [ranges[0][1] - ranges[0][0], ranges[1][1] - ranges[1][0], ranges[2][1] - ranges[2][0]]).max() / 2.0
    mid_x = np.mean(ranges[0])
    mid_y = np.mean(ranges[1])
    mid_z = np.mean(ranges[2])
    ax.set_xlim(mid_x - max_range, mid_x + max_range)
    ax.set_ylim(mid_y - max_range, mid_y + max_range)
    ax.set_zlim(mid_z - max_range, mid_z + max_range)


def make_pca_plot(X, use_3d=False, transient_frac=0.03, use_direct_projection=False, w=None, title='', dt=0.01, save_name=None, data_ranges=[[0, 0], [0, 0], [0, 0]],
                  flow_scale=None, bbox_lims = False):
    pca_model = None
    if isinstance(X, list):
        X_list = X
        X = np.vstack(X)
    else:
        X_list = [X]
    if not use_direct_projection:
        pca_model = PCA(n_components=X.shape[1])
        X_pca = pca_model.fit_transform(X)
    else:
        X_pca = X
    X_pca_list = np.split(X_pca, len(X_list), axis=0)
    fig = plt.figure(dpi=200, figsize=(4, 3))
    if not use_3d:
        plt.title(title, fontsize=14)
    T, N = X_pca_list[0].shape
    transient_t = int(T * transient_frac)
    t = np.linspace(0, T * dt, T)[transient_t:]
    if use_3d:
        ax = fig.add_subplot(projection='3d')
        ax.set_title(title)
        ax.set_xlabel('PC1 (A.U.)', fontsize=14)
        ax.set_ylabel('PC2 (A.U.)', fontsize=14)
        ax.set_zlabel('PC3 (A.U.)', fontsize=14)
        for axis in ax.xaxis, ax.yaxis, ax.zaxis:
            axis.labelpad = -10
            axis.set_ticks_position('none')
        ax.grid(False)
    for i, X_curr in enumerate(X_pca_list):
        X_pca_curr = X_curr[transient_t:]
        c = t / t[-1]
        if use_3d:
            set_axes_equal_range(X_pca_curr, data_ranges, ax, scale=0.8)

            points = X_pca_curr[:, :3].reshape(-1, 1, 3)
            segments = np.concatenate([points[:-1], points[1:]], axis=1)
            norm = plt.Normalize(c.min(), c.max())
            lc = Line3DCollection(segments, cmap='viridis', norm=norm, alpha=1)
            lc.set_array(c)
            lc.set_linewidth(1.75)
            p = ax.add_collection3d(lc)
            ax.scatter(X_pca_curr[:1, 0], X_pca_curr[:1, 1], X_pca_curr[:1, 2], c=c[:1], s=6)

            if w is not None:
                _, evecs, _ = get_eigenproperties(w)
                evecs_pca = pca_model.transform(np.stack([np.real(evecs[:, 0]), np.real(evecs[:, 1])]))
                origin = np.array([[0, 0, 0], [0, 0, 0], [0, 0, 0]])
                plt.quiver(*origin, evecs_pca[0, 0], evecs_pca[0, 1],evecs_pca[0, 2], color='red')
                plt.quiver(*origin, evecs_pca[1, 0], evecs_pca[1, 1],evecs_pca[1, 2], color='red')
            if i == len(X_list) - 1:
                ax.set_box_aspect((1, 1, 1))
                cbar = fig.colorbar(p, ax=ax, pad=0.05, shrink=0.9, ticks=[0,0.5, 1])
                cbar.set_label('t/T', rotation=90, size=14)
                cbar.ax.tick_params(labelsize=14)
                cbar.solids.set(alpha=1)
                return fig, ax, pca_model, data_ranges
        else:
            plt.scatter(X_pca_curr[:, 0], X_pca_curr[:, 1], c=t, s=1.5)
            if w is not None:
                xmin, xmax = np.min(X_pca_curr[:, :2]), np.max(X_pca_curr[:, :2])
                if bbox_lims:
                    xmin = - max(abs(xmin), abs(xmax))
                    xmax = -xmin
                scale = max(abs(xmin), abs(xmax)) * 4
                if flow_scale is not None:
                    scale = flow_scale
                # For flow field grid, set other PCs to their corresponding values at halfway through the recording
                pad = X_pca[T // 2, 2:]
                xy, flow = get_flow_field(w, pca_model, x_range=[xmin, xmax], pc_pad=pad, x_num=12)
                xy_dense, flow_dense = get_flow_field(w, pca_model, x_range=[xmin*1.25, xmax*1.25], y_range = [xmin, xmax*1.2], pc_pad=pad, x_num=100)
                flow_dense_mag = np.linalg.norm(flow_dense, axis = 1) ** (1/10)
                flow_dense_mag /= np.max(flow_dense_mag)
                plt.imshow(flow_dense_mag.reshape(-1, int(np.sqrt(flow_dense_mag.shape[0]))), cmap='gist_yarg', origin='lower', extent=[xmin*1.25, xmax*1.25, xmin, xmax*1.2], interpolation='bilinear', alpha = 0.8)
                plt.quiver(xy[:, 0], xy[:, 1], flow[:, 0], flow[:, 1], color='black', scale=scale, width=0.004, headwidth=4,
                           headlength=6)
                plt.xlim([xmin*1.25, xmax*1.25])
                plt.ylim([xmin, xmax*1.2])
            cb = plt.colorbar()
            tick_locator = plt.MaxNLocator(2)
            cb.locator = tick_locator
            cb.update_ticks()
            cb.set_label(label='t/T', size=14)
            plt.xlabel('PC1 (A.U.)', fontsize=14)
            plt.ylabel('PC2 (A.U.)', fontsize=14)
            ax = plt.gca()
            ax.spines[['right', 'top']].set_visible(False)
            plt.tight_layout()
    if save_name is not None:
                plt.savefig(os.path.join(fig_path, save_name), format="pdf", bbox_inches="tight")

def make_eigenvalue_plot(weights: dict, f_suffix = ''):
    plt.figure(dpi=150, figsize=(2.5,2.5))
    t = np.linspace(0, np.pi*2, 100)
    plt.plot(np.cos(t), np.sin(t), color = 'gray', linewidth = 1, alpha = 0.5)
    plt.xlabel(r'$\Re(\lambda)$', fontsize=16)
    plt.ylabel(r'$\Im(\lambda)$',fontsize=16)
    plt.gca().yaxis.labelpad=-10
    colors = ['red', 'blue']
    #zorders = [10, 0]
    zorders = [0,10] # flip to change order of blue and red
    idx = 0
    for type in weights:
        w = weights[type]
        evs = np.linalg.eigvals(w)
        evs_real = np.real(evs)
        evs_imag = np.imag(evs)
        arg = np.argsort(evs_real)
        np.random.shuffle(arg)
        evs_real = evs_real[arg]
        evs_imag = evs_imag[arg]
        plt.scatter(evs_real, evs_imag, label = type, color = colors[idx], s = 20, edgecolor='black', linewidth=0.1, alpha = 0.5, zorder = zorders[idx])
        idx += 1
    ax = plt.gca()
    ax.spines[['right', 'top']].set_visible(False)
    ax.set_aspect('equal')
    ax.xaxis.set_tick_params(labelsize=16)
    ax.yaxis.set_tick_params(labelsize=16)
    plt.legend(frameon=False, fontsize=11, labelspacing=0.25, handletextpad=-0.1, loc = 'upper left', bbox_to_anchor=(0.0, 1.0))
    ax.set_xticks([-1, 0, 1])
    ax.set_yticks([-1, 0, 1])
    plt.tight_layout()
    plt.savefig(os.path.join(fig_path,'eval_plot_{}.pdf'.format(f_suffix)))

# Plot some results
def plot_rslds_fit_results(rnn, rslds, q_elbos_lem, zhat_lem, xhat_lem, yhat_lem, dt = 0.01, f_suffix = '', activity_subplots = False, ff_chain = False):
    n_latent = xhat_lem.shape[1]
    plt.figure()
    plt.plot(q_elbos_lem[1:], label="Laplace-EM")
    plt.legend()
    plt.xlabel("Iteration")
    plt.ylabel("ELBO")
    plt.savefig(os.path.join(fig_path, 'rslds_learning_curve_{}.pdf'.format(f_suffix)), format="pdf",
                bbox_inches="tight")

    A = rslds.dynamics.As[0]
    A = (A - (1 - dt) * np.eye(len(A))) / dt # map from (J = (1-\alpha)*I_D + \alpha*B) scaling to B scaling (discrete to continuous)
    plt.figure()
    plt.title('gt connectivity')
    plt.imshow(rnn.w)
    plt.figure()
    plt.imshow(A)
    plt.title('rSLDS dynamics matrix')
    plt.xticks(np.arange(0, n_latent))
    plt.yticks(np.arange(0, n_latent))
    plt.xlabel('latent dimensions')
    plt.ylabel('latent dimensions')
    plt.savefig(os.path.join(fig_path, 'rslds_dynamics_matrix_{}.pdf'.format(f_suffix)), format="pdf",
                bbox_inches="tight")
    plt.colorbar()

    if activity_subplots:
        fig, ax = plt.subplots(2,1, dpi=150, figsize=(4,3), constrained_layout=True)
        ax[0].set_title('Ground truth', fontsize=14)
        ax[0].plot(rnn.t, rnn.y)
        ax[0].tick_params(labelbottom=False)
        lims_y = ax[0].get_ylim()
        ax[0].spines[['right', 'top']].set_visible(False)

        predicted = yhat_lem.T
        ax[1].set_title('Learned', fontsize=14)
        ax[1].plot(rnn.t, predicted.T)
        ax[1].set_xlabel('t', fontsize=14)
        ax[1].set_ylim(lims_y)
        ax[1].spines[['right', 'top']].set_visible(False)
        fig.supylabel('Neural Response (A.U.)', fontsize=13, y = 0.54)
        plt.savefig(os.path.join(fig_path, 'rslds_neural_response_gt_pred_{}.pdf'.format(f_suffix)))

        plt.figure()
        plt.title('Difference')
        plt.plot(rnn.t, predicted.T - rnn.y)
    else:
        plt.figure(dpi=150, figsize=(4,3))
        plt.title('Ground truth', fontsize=14)
        plt.plot(rnn.t, rnn.y)
        plt.xlabel('t', fontsize=14)
        plt.ylabel('Neural Response (A.U.)', fontsize=14)
        ax = plt.gca()
        lims_y = ax.get_ylim()
        ax.spines[['right', 'top']].set_visible(False)
        plt.tight_layout()
        plt.savefig(os.path.join(fig_path, 'rslds_neural_response_gt_{}.pdf'.format(f_suffix)))

        predicted = yhat_lem.T
        plt.figure(dpi=150, figsize=(4,3))
        plt.title('Learned', fontsize=14)
        plt.plot(rnn.t, predicted.T)
        plt.xlabel('t', fontsize=14)
        plt.ylabel('Neural Response (A.U.)', fontsize=14)
        plt.ylim(lims_y)
        ax = plt.gca()
        ax.spines[['right', 'top']].set_visible(False)
        plt.tight_layout()
        plt.savefig(os.path.join(fig_path, 'rslds_neural_response_predicted_{}.pdf'.format(f_suffix)))

    evs, evecs, tau_ev = get_eigenproperties(A)
    u_A, _, v_A = np.linalg.svd(A)

    flow_scales = [250, 1300]
    if ff_chain:
        flow_scales = [150, 525]
    make_pca_plot(xhat_lem, transient_frac=0.0, use_3d=False, w=A, title='Learned',
                  save_name='predicted_pca_{}.pdf'.format(f_suffix), flow_scale=flow_scales[0])
    evs_orig, evecs_orig, tau_orig = get_eigenproperties(rnn.w)

    make_pca_plot(rnn.x, transient_frac=0.0,use_3d=False, w=rnn.w, title='Ground Truth', save_name='ground_truth_pca_{}.pdf'.format(f_suffix),
                  flow_scale=flow_scales[1])

    plt.figure(dpi=150, figsize = (3,2.25))
    plt.plot(np.arange(1, n_latent + 1), tau_ev, '-o', color = 'blue')
    plt.ylabel('time constant ' + r'$\tau_n$', fontsize=13)
    plt.xticks(np.arange(1, n_latent + 1), fontsize=14)
    plt.yticks(fontsize=14)
    plt.xlabel('Dimension', fontsize=14)
    plt.title(f"Line attractor score: {np.log2(tau_ev[0] / tau_ev[1]):0.2f}", fontsize=12)
    ax = plt.gca()
    ylims_tau = ax.get_ylim()
    ax.spines[['right', 'top']].set_visible(False)
    plt.savefig(os.path.join(fig_path, 'rslds_time_constants_{}.pdf'.format(f_suffix)), format="pdf",
                bbox_inches="tight")
    plt.tight_layout()

    plt.figure(dpi=150, figsize = (3,2.25))
    if ff_chain: # correct small pseudospectral drift
        plt.plot(np.arange(1, n_latent + 1), np.ones(n_latent), '-o', color = 'red')
        plt.ylim([-20, 310])
    else:
        plt.plot(np.arange(1, n_latent + 1), tau_orig[:5], '-o', color='red')
        plt.ylim(ylims_tau)
    plt.ylabel('time constant ' + r'$\tau_n$', fontsize=13)
    plt.xticks(np.arange(1, n_latent + 1), fontsize=14)
    plt.yticks(fontsize=14)
    plt.xlabel('Dimension', fontsize=14)
    plt.title(f"Line attractor score: {np.log2(tau_orig[0] / tau_orig[1]):0.2f}", fontsize=12)
    ax = plt.gca()
    ax.spines[['right', 'top']].set_visible(False)
    plt.savefig(os.path.join(fig_path, 'rslds_time_constants_gt_{}.pdf'.format(f_suffix)), format="pdf",
                bbox_inches="tight")
    plt.tight_layout()
    plt.show()

def make_input_readout_plots(rnn, f_suffix='', subplot = False):
    if subplot:
        fig, ax = plt.subplots(2,1, figsize=(4,3))
        ax[0].plot(rnn.t, rnn.inp, color='gray', label = 'Input')
        ax[0].set_ylabel('Stimulus (A.U.)', fontsize=11)
        ax[0].legend(fontsize=11, frameon=False)
        ax[0].spines[['right', 'top']].set_visible(False)
        ax[0].tick_params(labelbottom=False)

        ax[1].plot(rnn.t, rnn.readout, label='Network', color='green')
        ax[1].plot(rnn.t, rnn.desired_out, label='Ideal', color='black')
        ax[1].set_xlabel('t', fontsize=14)
        ax[1].set_ylabel('Readout (A.U.)', fontsize=11)
        ax[1].legend(fontsize=11, frameon=False)
        ax[1].spines[['right', 'top']].set_visible(False)
        plt.tight_layout()
        plt.savefig(os.path.join(fig_path, 'input_readout_{}.pdf'.format(f_suffix)))
    else:
        plt.figure(dpi=150, figsize=(4,3))
        plt.title('Input', fontsize=14)
        plt.plot(rnn.t, rnn.inp, color='gray')
        plt.xlabel('t', fontsize=14)
        plt.ylabel('Stimulus (A.U.)', fontsize=14)
        ax = plt.gca()
        ax.spines[['right', 'top']].set_visible(False)
        plt.tight_layout()
        plt.savefig(os.path.join(fig_path, 'input_signal_{}.pdf'.format(f_suffix)))

        plt.figure(dpi=150, figsize=(4,3))
        plt.plot(rnn.t, rnn.readout, label='predicted output', color='green')
        plt.plot(rnn.t, rnn.desired_out, label='desired output', color='black')
        plt.xlabel('t', fontsize=14)
        plt.ylabel('Readout (A.U.)', fontsize=14)
        plt.legend(fontsize=9.5, frameon=False)
        ax = plt.gca()
        ax.spines[['right', 'top']].set_visible(False)
        plt.tight_layout()
        plt.savefig(os.path.join(fig_path, 'readout_{}.pdf'.format(f_suffix)))
```
