## Supplementary material for "Partial observation can induce mechanistic mismatches in data-constrained models of neural dynamics": Code: README.txt

Code for "Partial observation can induce mechanistic mismatches
in data-constrained models of neural dynamics".

To use the same simulation conditions (initial conditions, connectivity matrices, input noise, etc.) used for plots in the manuscript, we have provided an anonymized download link to the relevant files (too large to add directly to the supplementary files): 
https://www.dropbox.com/scl/fi/4306wf3tnvld9bzh963e9/additional_files_neurips_2024_mechanistic_mismatch.zip?rlkey=hyexf3cfwvdllmnzlx3z7dy45&st=ms364yqa&dl=0

Download and extract the zip such that it replaces the initially empty folders "saved_activity", "saved_conditions", and "saved_plot_data".

This code makes use of or directly contains code attributable to the following public repositories:

https://github.com/lindermanlab/ssm
https://github.com/rajanlab/CURBD
https://github.com/schnitzer-lab/CORNN-public

ssm must be first installed using the installation guide provided in the repository link. Relevant code snippets from the other two repositories have already been included. 
