## Supplementary material for "Partial observation can induce mechanistic mismatches in data-constrained models of neural dynamics": Code: RNN.py

```
from simulation_conditions import *

class RNN():
    def __init__(self, n_neuron, nt, alpha, v0, v1, n_obs, s_obs=0, g=1, phi = lambda x: x, conditions=None, integrate_sum = True, nonlinear_weights=False, network = 'gaussian',**kwargs):
        self.n_neuron = n_neuron # D
        self.nt = nt # timesteps
        self.alpha = alpha
        self.v0, self.v1 = v0, v1 # input signal scale, input noise scale
        self.phi, self.g = phi, g
        self.contribution_f = lambda z, w: self.g * self.phi(z) @ self.w.T
        self.nonlinear_weights = nonlinear_weights
        if nonlinear_weights: # for leaky-rate instead of leaky-current, as in CORNN
            self.contribution_f = lambda z, w: self.phi(self.g * z@ self.w.T)
        self.n_obs = n_obs # d
        self.s_obs = s_obs # observation noise scale. Set to 0 in all experiments
        self.t = alpha * np.arange(0, nt)
        self.x = np.zeros((nt, n_neuron))
        self.v = np.zeros((nt, n_neuron)) # inputs
        self.w_in, self.w_out, self.inp, self.readout = None, None, None, None
        if conditions is None: # if simulation conditions are not supplied, generate them
            conditions = generate_simulation_conditions(n_neuron, nt, n_obs, network=network,
                                                                            **kwargs)
        self.x[0, :] = conditions['ic']
        self.w = conditions['w']
        if conditions['U'] is not None and v0 > 0:
            self.set_inp_readout_weights(conditions['U'], integrate_sum)
        self.v += v1 * conditions['inp_noise']
        self.c = np.vstack((np.eye(n_obs), np.zeros((n_neuron - n_obs, n_obs))))
        self.obs_noise = conditions['obs_noise'] * s_obs
        self.forward()
    def set_inp_readout_weights(self, U, integrate_sum):
        integration_mode, self.inp, self.desired_out = generate_integration_input_output(U, self.nt, integrate_sum=integrate_sum)
        self.inp *= self.v0
        self.desired_out *= self.v0
        self.w_in = integration_mode
        self.w_out = integration_mode / self.n_neuron
        # rescale readouts for nonnormal integrator and line attractor. Adjusted to solve the task.
        if integrate_sum:
            self.w_out *= 0.7
        else:
            self.w_out /= self.n_neuron
        self.v += self.inp * self.w_in
        # ensure initialization is at 0 readout by projecting out the integration mode.
        self.x[0, :] -= np.dot(self.x[0, :], integration_mode) * integration_mode / (
                np.linalg.norm(integration_mode) ** 2)

    def forward(self):
        for i in range(self.nt - 1):
            self.x[i + 1, :] = self.x[i, :] + self.alpha * (self.contribution_f(self.x[i, :], self.w) - self.x[i, :] + self.v[i, :])
        if self.w_out is not None:
            self.readout = self.x @ self.w_out
        self.y = self.x @ self.c + self.obs_noise # observed neurons

def get_input_signal(nt):
    # random handpicked signal to integrate
    signal_to_integrate = np.zeros(nt)
    signal_to_integrate[1000:1200] = 1
    signal_to_integrate[1250:1450] = -1
    signal_to_integrate[800:950] = -2
    signal_to_integrate[1600:1800] = 3

    signal_to_integrate[2200:2400] = 0.5

    signal_to_integrate[2600:3000] = -np.linspace(0, 3, 400)
    signal_to_integrate[3500:4500] = 0.1

    signal_to_integrate = signal_to_integrate.reshape(-1, 1)
    signal_to_integrate = signal_to_integrate + np.random.standard_normal(nt).reshape(-1, 1) / 20
    return signal_to_integrate

def generate_integration_input_output(U, nt, alpha=0.01, integrate_sum=True):
    signal_to_integrate = get_input_signal(nt)
    output = np.cumsum(signal_to_integrate) * alpha # desired output

    if integrate_sum: # nonnormal integrator
        integration_mode = np.sum(U, axis=-1)
    else: # line attractor
        integration_mode = U[:, 0] * len(U)
    return integration_mode, signal_to_integrate, output
```
