## Supplementary material for "Partial observation can induce mechanistic mismatches in data-constrained models of neural dynamics": Code: RNN_fit.py

```
import copy
import ssm
from utils_cornn import solve_corrn
from utils_RNN import *


import math

import numpy as np
import numpy.random as npr
import numpy.linalg

import matplotlib.pyplot as plt
from matplotlib.gridspec import GridSpec

def fit_solution_direct(X_all, d, alpha, rho, phi=lambda x: x):
    '''
    :param X_all: (n_obs = d, T) activity
    :param d: observations
    :param alpha: discretization param
    :param rho: regularization
    :param phi: transfer function
    :return: fit
    '''
    X, Xs = X_all[:, :-1], X_all[:, 1:]
    X_phi = phi(X)
    first_mat = (Xs - (1 - alpha) * X) @ X_phi.T
    inv_arg = rho * np.eye(d) + alpha ** 2 * X_phi @ X_phi.T
    return alpha * first_mat @ np.linalg.pinv(inv_arg)

def fit_solution_implicit(B, Z, d, alpha, rho, input_noise, phi=lambda x: x, noise_limit=False):
    D = B.shape[0]
    if d < D:
        proj = np.hstack((np.eye(d), np.zeros((d, D - d))))
    else:
        proj = np.eye(d)
    if noise_limit:
        return get_learned_stationary(B, d)
    Z_phi = phi(Z)
    inv_arg = rho * np.eye(d) + alpha ** 2 * proj @ Z_phi @ Z_phi.T @ proj.T
    first_mat = proj @ (B @ Z_phi @ Z_phi.T + input_noise @ Z_phi.T) @ proj.T
    B_out = alpha ** 2 * first_mat @ np.linalg.pinv(inv_arg)
    return B_out

def fit_solution_grad_descent(X_all, dt=0.01, rho=0, lr=1e-2, max_iter=10000, phi=lambda x: x):
    d, T = X_all.shape
    A_hat = np.zeros((d, d))
    X, Xs = X_all[:, :-1], X_all[:, 1:]
    X_phi = phi(X)
    for i in range(max_iter):
        g = dt * ((1 - dt) * X + dt * A_hat @ X_phi - Xs) @ X_phi.T + rho * A_hat
        A_hat -= lr * g
    return A_hat

def fit_cornn(y, temporal_sample_every_n=1, alpha=0.01, inputs = None, solver_type = 'weighted'):
    y = y[::temporal_sample_every_n]
    r_in, r_out = y[:-1, :], y[1:, :]
    return solve_corrn(r_in, r_out, alph=alpha, num_iters=1000, l2=1e-5, u_in=inputs, threshold=1.0, solver_type=solver_type)

def trainMultiRegionRNN(activity, dtData=1, dtFactor=1, g=1.5, tauRNN=0.01,
                        tauWN=0.1, ampInWN=0.01, nRunTrain=2000,
                        nRunFree=10, P0=1.0,
                        nonLinearity=np.tanh,
                        nonLinearity_inv=np.arctanh,
                        resetPoints=None,
                        plotStatus=True, verbose=True,
                        regions=None):
    # FORCE, Adapted from https://github.com/rajanlab/CURBD
    r"""
    Trains a data-constrained multi-region RNN. The RNN can be used for,
    among other things, Current-Based Decomposition (CURBD).

    Parameters
    ----------

    activity: numpy.array
        N X T
    dtData: float
        time step (in s) of the training data
    dtFactor: float
        number of interpolation steps for RNN
    g: float
        instability (chaos); g<1=damped, g>1=chaotic
    tauRNN: float
        decay constant of RNN units
    tauWN: float
        decay constant on filtered white noise inputs
    ampInWN: float
        input amplitude of filtered white noise
    nRunTrain: int
        number of training runs
    nRunFree: int
        number of untrained runs at end
    P0: float
        learning rate
    nonLinearity: function
        inline function for nonLinearity
    resetPoints: list of int
        list of indeces into T. default to only set initial state at time 1.
    plotStatus: bool
        whether to plot data fits during training
    verbose: bool
        whether to print status updates
    regions: dict()
        keys are region names, values are np.array of indeces.
    """
    if dtData is None:
        print('dtData not specified. Defaulting to 1.');
        dtData = 1;
    if resetPoints is None:
        resetPoints = [0, ]
    if regions is None:
        regions = {}

    number_units = activity.shape[0]
    number_learn = activity.shape[0]

    dtRNN = dtData / float(dtFactor)
    nRunTot = nRunTrain + nRunFree

# set up everything for training

    learnList = npr.permutation(number_units)
    iTarget = learnList[:number_learn]
    iNonTarget = learnList[number_learn:]
    tData = dtData*np.arange(activity.shape[1])
    tRNN = np.arange(0, tData[-1] + dtRNN, dtRNN)

    ampWN = math.sqrt(tauWN/dtRNN)
    iWN = ampWN * npr.randn(number_units, len(tRNN))
    inputWN = np.ones((number_units, len(tRNN)))
    for tt in range(1, len(tRNN)):
        inputWN[:, tt] = iWN[:, tt] + (inputWN[:, tt - 1] - iWN[:, tt])*np.exp(- (dtRNN / tauWN))
    inputWN = ampInWN * inputWN

    # initialize directed interaction matrix J
    J = g * npr.randn(number_units, number_units) / math.sqrt(number_units)
    J0 = J.copy()

    # set up target training data
    Adata = activity.copy()
    # Adata = Adata/Adata.max()
    # Adata = np.minimum(Adata, 0.999)
    # Adata = np.maximum(Adata, -0.999)
    #Adata = nonLinearity(Adata)
    # get standard deviation of entire data
    stdData = np.std(Adata[iTarget, :])

    # get indices for each sample of model data
    iModelSample = numpy.zeros(len(tData), dtype=np.int32)
    for i in range(len(tData)):
        iModelSample[i] = (np.abs(tRNN - tData[i])).argmin()

    # initialize some others
    RNN = np.zeros((number_units, len(tRNN)))
    chi2s = []
    pVars = []

    # initialize learning update matrix (see Sussillo and Abbot, 2009)
    PJ = P0*np.eye(number_learn)

    if plotStatus is True:
        plt.rcParams.update({'font.size': 6})
        fig = plt.figure()
        fig.tight_layout()
        fig.subplots_adjust(hspace=0.4, wspace=0.4)
        gs = GridSpec(nrows=2, ncols=4)
    else:
        fig = None

    # start training
    # loop along training runs
    for nRun in range(0, nRunTot):
        H = Adata[:, 0, np.newaxis]
        RNN[:, 0, np.newaxis] = nonLinearity(H)

        # variables to track when to update the J matrix since the RNN and
        # data can have different dt values
        tLearn = 0  # keeps track of current time
        iLearn = 0  # keeps track of last data point learned
        chi2 = 0.0

        for tt in range(1, len(tRNN)):
            # update current learning time
            tLearn += dtRNN
            # check if the current index is a reset point. Typically this won't
            # be used, but it's an option for concatenating multi-trial data
            if tt in resetPoints:
                timepoint = math.floor(tt / dtFactor)
                H = Adata[:, timepoint]
            # compute next RNN step
            RNN[:, tt, np.newaxis] = nonLinearity(H)
            JR = (J.dot(RNN[:, tt]).reshape((number_units, 1)) +
                  inputWN[:, tt, np.newaxis])
            H = H + dtRNN*(-H + JR)/tauRNN
            # check if the RNN time coincides with a data point to update J
            if tLearn >= dtData:
                tLearn = 0
                #err = RNN[:, tt, np.newaxis] - Adata[:, iLearn, np.newaxis]
                err = RNN[:, tt, np.newaxis] - nonLinearity(Adata[:, iLearn, np.newaxis])
                iLearn = iLearn + 1
                # update chi2 using this error
                chi2 += np.mean(err ** 2)

                if nRun < nRunTrain:
                    r_slice = RNN[iTarget, tt].reshape(number_learn, 1)
                    k = PJ.dot(r_slice)
                    rPr = (r_slice).T.dot(k)[0, 0]
                    c = 1.0/(1.0 + rPr)
                    PJ = PJ - c*(k.dot(k.T))
                    J[:, iTarget.flatten()] = J[:, iTarget.reshape((number_units))] - c*np.outer(err.flatten(), k.flatten())

        rModelSample = RNN[iTarget, :][:, iModelSample]
        distance = np.linalg.norm(nonLinearity(Adata[iTarget, :]) - rModelSample)
        pVar = 1 - (distance / (math.sqrt(len(iTarget) * len(tData))
                    * stdData)) ** 2
        pVars.append(pVar)
        chi2s.append(chi2)
        if verbose:
            print('trial=%d pVar=%f chi2=%f' % (nRun, pVar, chi2))
            if nRun % 10 == 0:
                evs, _, _ = get_eigenproperties(J)
                print(nRun, evs[:2])
        if fig:
            fig.clear()
            ax = fig.add_subplot(gs[0, 0])
            ax.axis('off')
            ax.imshow(Adata[iTarget, :], aspect='auto')
            ax.set_title('real rates')

            ax = fig.add_subplot(gs[0, 1])
            ax.imshow(nonLinearity_inv(RNN), aspect='auto')
            ax.set_title('model rates')
            ax.axis('off')

            ax = fig.add_subplot(gs[1, 0])
            ax.plot(pVars)
            ax.set_ylabel('pVar')

            ax = fig.add_subplot(gs[1, 1])
            ax.plot(chi2s)
            ax.set_ylabel('chi2s')

            ax = fig.add_subplot(gs[:, 2:4])
            idx = npr.choice(range(len(iTarget)))
            idx=8
            ax.plot(tRNN, nonLinearity_inv(RNN[iTarget[idx], :]))
            ax.plot(tData, Adata[iTarget[idx], :])
            ax.set_title(nRun)
            fig.show()
            plt.pause(0.05)

    out_params = {}
    out_params['dtFactor'] = dtFactor
    out_params['number_units'] = number_units
    out_params['g'] = g
    out_params['P0'] = P0
    out_params['tauRNN'] = tauRNN
    out_params['tauWN'] = tauWN
    out_params['ampInWN'] = ampInWN
    out_params['nRunTot'] = nRunTot
    out_params['nRunTrain'] = nRunTrain
    out_params['nRunFree'] = nRunFree
    out_params['nonLinearity'] = nonLinearity
    out_params['resetPoints'] = resetPoints

    out = {}
    out['regions'] = regions
    out['RNN'] = RNN
    out['tRNN'] = tRNN
    out['dtRNN'] = dtRNN
    out['Adata'] = Adata
    out['tData'] = tData
    out['dtData'] = dtData
    out['J'] = J
    out['J0'] = J0
    out['chi2s'] = chi2s
    out['pVars'] = pVars
    out['stdData'] = stdData
    out['inputWN'] = inputWN
    out['iTarget'] = iTarget
    out['iNonTarget'] = iNonTarget
    out['params'] = out_params
    return out
def fit_rslds(y, n_obs, k=1, n_latent=5, temporal_sample_every_n=1, n_iters=200, input=None):
    # Fit an rSLDS with its default initialization, using Laplace-EM with a structured variational posterior
    M = 0
    if input is not None:
        M = input.shape[-1]
    rslds = ssm.SLDS(n_obs, k, n_latent, M=M,
                     transitions="recurrent_only",
                     dynamics="diagonal_gaussian",
                     emissions='gaussian_orthog',
                     single_subspace=True)
    y = y[::temporal_sample_every_n]
    rslds.initialize(y, inputs=input)
    q_elbos_lem, q_lem = rslds.fit(y, inputs=input, method="laplace_em",
                                   variational_posterior="structured_meanfield",
                                   initialize=False, num_iters=n_iters, alpha=0.0)
    xhat_lem = q_lem.mean_continuous_states[0]
    zhat_lem = rslds.most_likely_states(xhat_lem, y)

    yhat_lem = rslds.smooth(xhat_lem, y, input=input)

    # store rslds
    rslds_lem = copy.deepcopy(rslds)

    return rslds, q_elbos_lem, zhat_lem, xhat_lem, yhat_lem
```
