## Supplementary material for "Partial observation can induce mechanistic mismatches in data-constrained models of neural dynamics": Code: simulation_conditions.py

```
import scipy
from scipy.stats import ortho_group
from numpy.random import default_rng
import numpy as np
from utils_RNN import *

rng = default_rng()

def generate_simulation_conditions(n_neuron, nt, n_obs, x0_scale=1, **kwargs):
    '''

    :param n_neuron: teacher size (D)
    :param nt: timesteps
    :param n_obs: observation/student size (d)
    :param x0_scale: initial conditions (ICs) scale
    :param kwargs: teacher network parameters
    :return: ICs, teacher weights, input noise, observation noise, transform
    '''
    x0 = rng.standard_normal(n_neuron) * x0_scale
    w, U = get_W(n_neuron, **kwargs)
    inp_noise = rng.standard_normal((nt, n_neuron))
    obs_noise = rng.standard_normal((nt, n_obs))
    return {'ic': x0, 'w': w, 'inp_noise': inp_noise, 'obs_noise': obs_noise, 'U': U}

def generate_jordan_block(n_neuron):
    J = np.zeros((n_neuron, n_neuron))
    for i in range(n_neuron - 1):
        J[i][i + 1] = 1
    return J

def henrici_nonnormality(A):
    evals = np.linalg.eigvals(A)
    norm = np.linalg.norm(A)
    return np.sqrt(np.abs(norm ** 2 - np.sum(np.abs(evals) ** 2))) / norm

def get_delay_line(n_neuron, diagonal_comp=0, scale=1, skip_interval=0,
                   skip_scale=0.5):
    delay = np.zeros((n_neuron, n_neuron))
    if skip_interval > 0:
        delay[0][1 + skip_interval::skip_interval] = skip_scale
    delay += generate_jordan_block(n_neuron) * scale
    out = delay + np.eye(n_neuron) * diagonal_comp
    return out

def get_triangular(n_neuron, scale=3, diagonal_comp=0, use_schur = False):
    Q = np.random.normal(0, scale / np.sqrt(n_neuron), (n_neuron, n_neuron))
    if not use_schur:
        Q[np.tril_indices(n_neuron)] = 0
        out = Q + np.eye(n_neuron) * diagonal_comp
    else:
        T, Z = scipy.linalg.schur(Q, output='complex')
        diag_idxes = np.diag_indices(n_neuron)
        T[diag_idxes] = T[diag_idxes] / np.max(np.abs(T[diag_idxes])) * diagonal_comp
        out = np.real(Z @ T @ Z.T.conj())
    return out

def get_gaussian_network(n_neuron, scale=0.8, self_loops=True):
    Q = np.random.normal(0, scale / np.sqrt(n_neuron), (n_neuron, n_neuron))
    if not self_loops:
        for i in range(n_neuron):
            Q[i][i] = 0.0
    return Q

def get_eigenproperties(A, get_singular_values = False):
    evs, evecs = np.linalg.eig(A)
    idx = np.flip(np.argsort(np.real(evs)))
    evs = evs[idx]
    tau_ev = np.abs(1/(1 - np.real((evs))))
    if get_singular_values:
        _, s, _ = np.linalg.svd(A)
        return evs, evecs, tau_ev, s
    return evs, evecs, tau_ev

def get_symmetric_network(n_neuron, scale=0.6, self_loops=True, line_attractor=False):
    Q = np.random.normal(0, scale / np.sqrt(n_neuron), (n_neuron, n_neuron))
    if not self_loops:
        for i in range(n_neuron):
            Q[i][i] = 0
    out = (Q + Q.T) / 2
    if line_attractor:
        _, evecs, _ = get_eigenproperties(out)
        evs = np.linspace(-0.4, 0.4, n_neuron)[::-1]
        evs[0] = 0.999
        out = evecs @ np.diag(evs) @ evecs.T
    return out
    # return out

def get_low_rank_random(n_neuron, rank=2, scale=1):
    rand = np.random.standard_normal((n_neuron, 2 * rank))
    rand = rand / np.linalg.norm(rand, axis=0)
    Mm = rand[:, :rank]
    Nm = rand[:, rank:2 * rank]
    out = Mm @ Nm.T * scale
    return out

def get_low_rank_nonnormal_osc(n_neuron, shift_scale=12):
    Sigma = np.zeros((2, 2))
    Sigma[0, 0] = 0.5
    Sigma[1, 1] = 0.5
    Sigma[0, 1] = -0.5
    Sigma[1, 0] = 0.5
    M_mat = np.random.randn(n_neuron, 2) # relaxed the orthogonality constraint
    M_pinv = np.linalg.pinv(M_mat)
    I = np.eye(n_neuron)
    U_shift = np.random.normal(0, shift_scale / np.sqrt(n_neuron), (2, n_neuron))
    M_pinv += U_shift @ (I - M_mat @ M_pinv)
    N_mat_T = Sigma @ M_pinv
    out = M_mat @ N_mat_T
    return out
def get_low_rank_nonnormal(n_neuron, rank=2, scale=1, use_overlap_mat=False):

    # obtain orthogonal columns via tall and thin random matrix. More efficient than ortho.rvs -> square matrix
    rand = np.random.standard_normal((n_neuron, 2 * rank))
    Q, R = np.linalg.qr(rand, mode='reduced')
    L = np.sign(np.diag(R))  # for QR via Householder reflections -> Haar distribution fix
    Q = Q * L[None, :]

    Mm = Q[:, :rank]
    Nm = Q[:, rank:2 * rank]

    gam_sq = n_neuron * scale / np.sqrt(rank)
    Mm *= np.sqrt(gam_sq)
    Nm *= np.sqrt(gam_sq)
    out = Mm @ Nm.T

    return out

def get_specified_rank_normal(n_neuron, max_eval=0.999, min_eval=0.2, rank=2, diagonal=False):
    U = ortho_group.rvs(n_neuron)
    if diagonal:
        U = np.eye(n_neuron)
    evals = np.ones(n_neuron) * min_eval
    evals[0] = max_eval
    return U[:, :rank] @ np.diag(evals[:rank]) @ U[:, :rank].T

def get_W(n_neuron, network='gaussian', ortho_transform=False, perm_transform=False, similarity_transform=False,
          **kwargs):
    out = None
    if network == 'gaussian':
        out = get_gaussian_network(n_neuron, **kwargs)
    elif network == 'symmetric':
        out = get_symmetric_network(n_neuron, **kwargs)
    elif network == 'triangular':
        out = get_triangular(n_neuron, **kwargs)
    elif network == 'delay':
        out = get_delay_line(n_neuron, **kwargs)
    elif network == 'lr_nn':
        out = get_low_rank_nonnormal(n_neuron, **kwargs)
    elif network == 'lr_nn_osc':
        out = get_low_rank_nonnormal_osc(n_neuron, **kwargs)
    elif network == 'lr_n':
        out = get_specified_rank_normal(n_neuron, **kwargs)
    elif network == 'lr_r':
        out = get_low_rank_random(n_neuron, **kwargs)
    else:
        print(network)
        raise ValueError('unknown network type')
    U = None
    if ortho_transform:
        U = ortho_group.rvs(n_neuron)
        out = U @ out @ U.T
    if perm_transform:
        I = np.eye(n_neuron)
        p = np.random.permutation(n_neuron)
        U = I[p]
        out = U @ out @ U.T
    if similarity_transform:
        U = np.random.normal(0, 1 / np.sqrt(n_neuron), (n_neuron, n_neuron))
        U = U / np.linalg.norm(U, axis=0, keepdims=True)
        out = U @ out @ np.linalg.inv(U)
    return out, U
```
