## Supplementary material for "Partial observation can induce mechanistic mismatches in data-constrained models of neural dynamics": Code: sweep_plots.py

```
import numpy as np
import pickle as pk
import os

from matplotlib.colors import LinearSegmentedColormap
from scipy.stats import sem
import matplotlib.pyplot as plt

from RNN import RNN
from RNN_fit import fit_solution_direct
from simulation_conditions import generate_simulation_conditions, get_eigenproperties, get_W
from utils_RNN import get_stationary_cov, get_stationary_fit, get_subsampling_curves, henrici_nonnormality

# Make a folder to store figures
fig_path = './figures'
if not os.path.isdir(fig_path):
    os.makedirs(fig_path)
plot_data_path = './saved_plot_data'
print(f"Storing figures in {fig_path}")

def make_low_rank_finite_time_plots(n_obs=25, n_neuron=500, scale = 1, r_max=5, nt_finite=30000, v1=0.2, rho=0.001, alpha=0.01, trials = 20):
    evs_finite_all, evs_limit_all = [], []
    f_prefix = 'lr_nn_ft_plots'
    lr_ft_data_path = os.path.join(plot_data_path, 'low_rank_ft_effects')
    if len(os.listdir(lr_ft_data_path)) == 0:
        print('Could not find saved data; generating from scratch.')
        for r in range(1, r_max+1):
            evs_finite_curr, evs_limit_curr = [], []
            for n in range(trials):
                conditions = generate_simulation_conditions(n_neuron, nt_finite, n_obs, network='lr_nn', rank=r, scale=scale)
                rnn = RNN(n_neuron, nt_finite, alpha, 0.0, v1, n_obs, 0, conditions = conditions)
                B = conditions['w']
                cov = get_stationary_cov(B)
                A_limit = get_stationary_fit(B, n_obs, cov, ff_chain=False)
                A_finite = fit_solution_direct(rnn.y.T, n_obs, alpha, rho, phi=lambda x: x)
                evs_limit, _, _ = get_eigenproperties(A_limit)
                evs_finite, _, _ = get_eigenproperties(A_finite)
                evs_limit_curr.append(np.real(evs_limit[:10]))
                evs_finite_curr.append(np.real(evs_finite[:10]))
            evs_finite_all.append(evs_finite_curr)
            evs_limit_all.append(evs_limit_curr)
    else:
        evs_finite_all, evs_limit_all = pk.load(open(os.path.join(lr_ft_data_path, '{}_data_{}.pk'.format(f_prefix,scale)), 'rb'))
    evs_finite_all, evs_limit_all = np.array(evs_finite_all), np.array(evs_limit_all) #(r_max, trials, 10)
    evs_finite_mean, evs_limit_mean = np.mean(evs_finite_all, axis=1), np.mean(evs_limit_all, axis=1)
    evs_finite_sem, evs_limit_sem = sem(evs_finite_all, axis = 1), sem(evs_limit_all, axis=1)
    x_axis = np.arange(1, 11)
    x_label = 'Dimension'
    fig, ax = plt.subplots(2,2, dpi=150, figsize=(5.25,3), sharex=True, sharey=True, layout='constrained')
    colors = ['cadetblue', 'chocolate']
    for i in range(1, r_max):
        r, c = (i-1) // 2, (i-1) % 2
        label = None
        if i == 1:
            label = 'Finite time'
        ax[r,c].plot (x_axis, evs_finite_mean[i], label=label, color = colors[0], linestyle='--', marker= 'o', markerfacecolor='none', markersize=6, linewidth=0.75, markeredgewidth=0.75)
        ax[r,c].errorbar(x_axis, evs_finite_mean[i], yerr=evs_finite_sem[i], color=colors[0],ls='none', capsize=1.5, elinewidth=0.8, markeredgewidth=0.8)
        ax[r,c].spines[['right', 'top']].set_visible(False)
        ax[r,c].set_title('Rank {}'.format(i+1), fontsize=12)
    for i in range(1, r_max):
        r, c = (i - 1) // 2, (i - 1) % 2
        label = None
        if i == 1:
            label = r'$T \rightarrow \infty$'
        ax[r, c].plot(x_axis, evs_limit_mean[i], label = label, color = colors[1], markersize= 6, marker='x', linewidth=0.75, markeredgewidth=0.75)
        ax[r, c].errorbar(x_axis, evs_limit_mean[i], yerr=evs_limit_sem[i], color=colors[1], ls='none', capsize=1.5, elinewidth=0.8, markeredgewidth=0.8)

    fig.supxlabel(x_label, fontsize=14, x = 0.4)
    fig.supylabel(r'$\Re(\hat{\lambda})$', fontsize=14)
    lines_labels = [ax.get_legend_handles_labels() for ax in fig.axes]
    lines, labels = [sum(lol, []) for lol in zip(*lines_labels)]
    fig.legend(lines, labels, fontsize=12, frameon=False, loc="outside right center")
    plt.savefig(os.path.join(fig_path, '{}.pdf'.format(f_prefix)))

def make_ff_chain_plots(n_obs = 25):
    f_prefix = 'ff_chain'
    ff_chain_data_path = os.path.join(plot_data_path, 'ff_chain')
    n_neurons = np.arange(100, 1100, 100) # fig a
    evs_all, las_all = [], []
    if len(os.listdir(ff_chain_data_path)) == 0:
        print('Could not find saved data; generating from scratch.')
        #figs b,c
        n_neurons_large = (np.logspace(0, 2.30103, num=10, endpoint=True) * 25).astype(int)
        for n_neuron in n_neurons_large:
            print(n_neuron)
            B, _  = get_W(n_neuron, network='delay')
            cov = get_stationary_cov(B)
            A = get_stationary_fit(B, n_obs, cov, ff_chain=True)
            evs, _, _ = get_eigenproperties(A)
            evs_all.append(np.real(evs[:5]))

        for n_neuron in n_neurons:
            print(n_neuron)
            B, _, evs, las, _, _, _, _ = get_subsampling_curves(n_neuron,ff_chain=True,detailed=False, interval=1, network = 'delay')
            evs_all.append(evs)
            las_all.append(las)
    else:
        n_neurons_large, evs_increasing_D = pk.load(open(os.path.join(ff_chain_data_path, '{}_increasing_D_data_{}.pk'.format(f_prefix,n_obs)), 'rb'))
        evs_all, las_all = pk.load(open(os.path.join(ff_chain_data_path, '{}_subsampling_curves_data.pk'.format(f_prefix)), 'rb'))

    x_label = 'Subsampling fraction'
    colors = [(75/256, 0, 130/256), (1, 0.5, .25)]  # first color is purple, last is orangish
    cm = LinearSegmentedColormap.from_list(
        "Custom", colors, N=10)
    plt.figure(dpi=150, figsize=(4, 3))
    colors = ['#1f77b4', '#ff7f0e']
    top_evals_at_sub_frac = np.array([evs[:2] for evs in evs_increasing_D])
    timescales = 1/(1 - top_evals_at_sub_frac)
    plt.plot(n_neurons_large, timescales[:, 0], label = r'$\hat{\tau}_1$', color = colors[0])
    plt.scatter(n_neurons_large, timescales[:, 0], color = colors[0])

    plt.plot(n_neurons_large, timescales[:, 1], label = r'$\hat{\tau}_2$', color = colors[1])
    plt.scatter(n_neurons_large, timescales[:, 1], color = colors[1])
    plt.xscale('log')

    plt.xlabel('D (Teacher size)', fontsize = 14)
    plt.ylabel('Time constant '+r'$\hat{\tau}$', fontsize=14)
    plt.legend(frameon = False, fontsize=14)
    plt.tight_layout()
    plt.savefig(os.path.join(fig_path, '{}_timescales_vs_D_d_25.pdf'.format(f_prefix)))

    plt.figure(dpi=150, figsize=(4,3))
    plt.plot(n_neurons_large, top_evals_at_sub_frac[:, 0], label = r'$\hat{\lambda}_1$', color = colors[0])
    plt.scatter(n_neurons_large, top_evals_at_sub_frac[:, 0], color = colors[0])

    plt.plot(n_neurons_large, top_evals_at_sub_frac[:, 1], label = r'$\hat{\lambda}_2$', color = colors[1])
    plt.scatter(n_neurons_large, top_evals_at_sub_frac[:, 1], color = colors[1])
    plt.xscale('log')

    plt.xlabel('D (Teacher size)', fontsize = 14)
    plt.ylabel(r'$\Re(\hat{\lambda})$', fontsize=14)
    plt.legend(frameon = False, fontsize=14)
    plt.tight_layout()
    plt.savefig(os.path.join(fig_path, '{}_evals_vs_D_d_25.pdf'.format(f_prefix)))

    plt.figure(dpi=150, figsize=(4,3))
    plt.xscale('log', base=10)
    for i in range(len(n_neurons)):
        x_axis = np.arange(1, n_neurons[i]+1) / n_neurons[i]
        plt.plot(x_axis[1:], las_all[i][1:], label=r'$D={}$'.format(n_neurons[i]), color=cm(i/10))
    plt.xlabel(x_label, fontsize=14)
    plt.ylabel('Line attractor score', fontsize=14)
    plt.legend(frameon=False, fontsize=9)
    plt.tight_layout()
    plt.savefig(os.path.join(fig_path, '{}_las_vs_subsampling_at_D.pdf'.format(f_prefix)))


def make_subsampling_plots(n_neuron=500, interval=1, f_prefix='', show_top_n=2, trials = 10, **kwargs):
    subsampling_data_path = os.path.join(plot_data_path, 'subsampling_sweeps')
    saved_data_path = os.path.join(subsampling_data_path, '{}_subsampling_curves_data.pk'.format(f_prefix))
    h_vals_all, evs_all, las_all = [], [], []
    svs_all, uvecs_olap_all, vvecs_olap_all, evecs_olap_all = [], [], [], []
    evs_B_all, h_B_all, svs_B_all = [], [], []
    if not os.path.isfile(saved_data_path):
        print('Could not find saved data; generating from scratch.')
        for i in range(trials):
            B, h_vals, evs, las, svs, uvecs_overlap, vvecs_overlap, evecs_overlap = get_subsampling_curves(n_neuron, interval=interval, **kwargs)
            h_vals_all.append(h_vals); evs_all.append(evs); las_all.append(las)
            svs_all.append(svs); uvecs_olap_all.append(uvecs_overlap); vvecs_olap_all.append(vvecs_overlap); evecs_olap_all.append(evecs_overlap)
            evs_B, _, _, svs_B = get_eigenproperties(B, get_singular_values=True)
            evs_B = np.real(evs_B)
            h_B = henrici_nonnormality(B)
            evs_B_all.append(evs_B); h_B_all.append(h_B); svs_B_all.append(svs_B)
    else:
        h_vals_all, evs_all, las_all, svs_all, uvecs_olap_all, vvecs_olap_all, evecs_olap_all, evs_B_all, h_B_all, svs_B_all = pk.load(open(saved_data_path, 'rb'))
    h_vals_all, evs_all, las_all = np.array(h_vals_all), np.array(evs_all), np.array(las_all)

    svs_all, uvecs_olap_all, vvecs_olap_all, evecs_olap_all = np.array(svs_all), np.array(uvecs_olap_all), np.array(vvecs_olap_all), np.array(evecs_olap_all)
    evs_B_all, h_B_all, svs_B_all = np.array(evs_B_all), np.array(h_B_all), np.array(svs_B_all)

    h_vals_mean, evs_mean, las_mean = np.mean(h_vals_all, axis = 0), np.mean(evs_all, axis = 0), np.mean(las_all, axis = 0)
    svs_mean, uvecs_olap_mean, vvecs_olap_mean, evecs_olap_mean = np.mean(svs_all, axis = 0), np.mean(uvecs_olap_all, axis = 0), np.mean(vvecs_olap_all, axis = 0), np.mean(evecs_olap_all, axis = 0)
    evs_B_mean, h_B_mean, svs_B_mean = np.mean(evs_B_all, axis = 0), np.mean(h_B_all, axis = 0), np.mean(svs_B_all, axis = 0)

    h_vals_sem, evs_sem, las_sem = sem(h_vals_all, axis = 0), sem(evs_all, axis = 0), sem(las_all, axis = 0)
    svs_sem, uvecs_olap_sem, vvecs_olap_sem, evecs_olap_sem = sem(svs_all, axis = 0), sem(uvecs_olap_all, axis = 0), sem(vvecs_olap_all, axis = 0), sem(evecs_olap_all, axis = 0)
    evs_B_sem, h_B_sem, svs_B_sem = sem(evs_B_all, axis = 0), sem(h_B_all, axis = 0), sem(svs_B_all, axis=0)

    x_axis = np.arange(1, len(h_vals_mean) + 1)/len(h_vals_mean)
    x_label = 'Subsampling fraction'

    plt.figure(dpi=150, figsize=(4,3))
    plt.plot(x_axis[1:], las_mean[1:])
    plt.fill_between(x_axis[1:], las_mean[1:] - las_sem[1:], las_mean[1:] + las_sem[1:], color='gray', alpha=0.2)
    plt.ylabel('Line attractor score', fontsize=14)
    plt.xlabel(x_label, fontsize=14)
    plt.axhline(y=1.0, color = 'green')
    plt.xscale('log')
    ax = plt.gca()
    ax.spines[['right', 'top']].set_visible(False)
    plt.ylim([-0.3, 7])
    plt.tight_layout()
    plt.savefig(os.path.join(fig_path, '{}_las_curve.pdf'.format(f_prefix)))

    plt.figure(dpi=150, figsize=(4,3))
    plt.plot(x_axis, svs_mean[:, :show_top_n], label =  [r'$\hat{\sigma}_%s$' % (i+1) for i in range(show_top_n)])
    for i in range(show_top_n):
        plt.fill_between(x_axis, svs_mean[:, i] - svs_sem[:, i], svs_mean[:, i] + svs_sem[:, i], color='gray', alpha=0.2)
    for i in range(show_top_n):
        plt.scatter([x_axis[-1]], [svs_B_mean[i]], label = r'$\sigma_{}$'.format(i+1), marker='x')
    plt.xlabel(x_label, fontsize=14)
    plt.ylabel('Singular value', fontsize=14)
    plt.xscale('log')
    ax = plt.gca()
    ax.spines[['right', 'top']].set_visible(False)
    plt.tight_layout()
    plt.legend(frameon=False)
    plt.savefig(os.path.join(fig_path, '{}_top_svs_curve.pdf'.format(f_prefix)))
    # plt.yscale('log')
    # plt.ylim([0.5, 20])

    olap_colors = ['teal', 'darkmagenta']
    plt.figure(dpi=150, figsize=(4,3))
    spurious_overlap_scale = 1 / np.sqrt(np.arange(1, len(h_vals_mean) + 1))
    # spurious_overlap_scale = spurious_overlap_scale * correction_factor
    plt.plot(x_axis, uvecs_olap_mean, label=r'$O([\mathbf{u}_{1}]_{1:d}, \hat{\mathbf{u}}_1)$', color = olap_colors[0])
    plt.fill_between(x_axis, uvecs_olap_mean - uvecs_olap_sem, uvecs_olap_mean + uvecs_olap_sem, color='gray', alpha=0.2)
    plt.plot(x_axis, vvecs_olap_mean, label=r'$O([\mathbf{v}_{1}]_{1:d}, \hat{\mathbf{v}}_1)$', color = olap_colors[1])
    plt.fill_between(x_axis, vvecs_olap_mean - vvecs_olap_sem, vvecs_olap_mean + vvecs_olap_sem, color='gray', alpha=0.2)
    plt.plot(x_axis, spurious_overlap_scale, label='spurious' + r'$\sim \frac{1}{\sqrt{d}}$',
             color='red')
    plt.xscale('log')
    plt.xlabel(x_label, fontsize=14)
    plt.ylabel('Overlap', fontsize=14)
    plt.legend(frameon=False, fontsize=10, loc='center right')
    ax = plt.gca()
    ax.spines[['right', 'top']].set_visible(False)
    plt.tight_layout()
    plt.savefig(os.path.join(fig_path, '{}_s_overlap_curve.pdf'.format(f_prefix)))

if __name__ == "__main__":
    make_low_rank_finite_time_plots(trials=20, n_neuron=500, n_obs=25, nt_finite=30000)
    make_ff_chain_plots()

    # load from file
    make_subsampling_plots(f_prefix='nn_integrator_0_5')

    # generate from scratch (uncomment)
    # make_subsampling_plots(500, f_prefix='', network='delay', skip_scale=0.25, skip_interval=1, ortho_transform=True, trials = 20)

    # load from file
    make_subsampling_plots(f_prefix='rank_1_gamma_0_4')

    # generate from scratch (uncomment)
    # make_subsampling_plots(500, f_prefix='', network='lr_nn', scale = 0.1, rank = 1, trials = 20)

    # etc
    plt.show()
```
