## Supplementary material for "Partial observation can induce mechanistic mismatches in data-constrained models of neural dynamics": Code: symmetric_teacher_plots.py

```
import os
import pickle as pk

import numpy as np
from matplotlib import pyplot as plt

from make_RNN_plots import  create_fit_plots
from simulation_conditions import generate_simulation_conditions

# Make a folder to store figures
fig_path = './figures'
if not os.path.isdir(fig_path):
    os.makedirs(fig_path)
conditions_path = './saved_conditions'
activity_path = './saved_activity'
print(f"Storing figures in {fig_path}")

if __name__ == "__main__":
    load_from_file = True
    if len(os.listdir(conditions_path)) == 0:
        print('No condition files found.')
        load_from_file = False

    '''No line attractor'''
    f_name = 'symmetric_no_la'
    if load_from_file:
        conditions = pk.load(open(os.path.join(conditions_path, f_name+'.pk'), 'rb'))
    else:
        conditions = generate_simulation_conditions(500, 30000, 25, network='symmetric', line_attractor=False)
    create_fit_plots(500, 30000, 25, v1=0.2, rho=0.001, fit_method='direct', conditions=conditions,
                     phi=lambda x: x, use_direct_projection=False, f_suffix=f_name,
                     nonlinear_weights=False, plot_time=4000, data=None, plot_flowfields=True)

    create_fit_plots(500, 30000, 25, v1=0.2, rho=0.001, fit_method='long_time_limit', conditions=conditions,
                     phi=lambda x: x, use_direct_projection=False, f_suffix=f_name+'_noise_lim',
                     nonlinear_weights=False, plot_time=4000, data=None, plot_flowfields=True)

    '''line attractor'''
    f_name = 'symmetric_la'
    if load_from_file:
        conditions = pk.load(open(os.path.join(conditions_path, f_name + '.pk'), 'rb'))
    else:
        conditions = generate_simulation_conditions(500, 30000, 25, network='symmetric', line_attractor=True)
    create_fit_plots(500, 30000, 25, v1=0.2, rho=0.001, fit_method='direct', conditions=conditions,
                     phi=lambda x: x, use_direct_projection=False, f_suffix=f_name,
                     nonlinear_weights=False, plot_time=4000, data=None, plot_flowfields=True)

    create_fit_plots(500, 30000, 25, v1=0.2, rho=0.001, fit_method='long_time_limit', conditions=conditions,
                     phi=lambda x: x, use_direct_projection=False, f_suffix=f_name + '_noise_lim',
                     nonlinear_weights=False, plot_time=4000, data=None, plot_flowfields=True)
    plt.show()
```
