## Supplementary material for "Partial observation can induce mechanistic mismatches in data-constrained models of neural dynamics": Code: utils_cornn.py

```
#!/usr/bin/env python3
# -*- coding: utf-8 -*-
"""
Created on Fri Jun 10 14:48:40 2022

@author: dinc
source: https://github.com/schnitzer-lab/CORNN-public
"""
import numpy as np
from scipy.linalg import solve
import re
import time as time_now
from numpy.linalg import inv
from scipy.stats import pearsonr
import multiprocessing
from joblib import Parallel, delayed
import torch
import torch.nn as nn
import torch.optim as optim

def solve_corrn(r_in,r_out,u_in = None, alph = 0.1, 
                l2 = 1e-4, threshold = 1, initialize_fp = 1,
                verbose = 0,check_convergence = 0,
                num_iters = 30,gnd = None,solver_type = 'weighted'):
    if verbose:
        temp = time_now.localtime()
        current_time = time_now.strftime("%H:%M:%S", temp)
        print('%s: Running CoRNN solver.' %(current_time))
    start_time = time_now.perf_counter()
    
    T_data = r_in.shape[0]
    n_rec = r_in.shape[1]
    if u_in is not None:
        n_in = u_in.shape[1]
    else:
        n_in = 0
    
    # Get the inputs x and targets d
    if n_in >0:
        x = np.c_[r_in,u_in];
    else:
        x = r_in.copy()
    d = (r_out - (1-alph)*r_in)/alph
    d[d<=-1+1e-6] = -1 + 1e-6;
    d[d>=1-1e-6]  = 1-1e-6;
    
    if verbose == 2:
        temp = time_now.localtime()
        current_time = time_now.strftime("%H:%M:%S", temp)
        print('\t %s: Performing initial inverse operations' %(current_time))
    
    # Scale the regularization and compute the fixed point
    l2  = l2*T_data;
        
    
    A = x.T @ x + (l2) * np.diag(np.ones(x.shape[1]));
    n_tot = n_rec + n_in
    
    Ainv = inv(A);
    Xp   = Ainv @ x.T;
    Xm  = Xp @ x; 
    z    = np.arctanh(d);
    theta_fp = Xp @ z;
    if initialize_fp:
        theta = theta_fp;
    else:
        theta = np.random.normal(0,1/np.sqrt(n_rec),n_rec * n_tot).reshape(n_tot,n_rec)
    
    if verbose == 2:
        temp = time_now.localtime()
        current_time = time_now.strftime("%H:%M:%S", temp)
        print('\t %s: Starting iterations' %(current_time))
        
    if ( (verbose == 2) & (gnd is not None) ) :
        prd = (theta[:n_rec,:].T).flatten();
        p_cor = pearsonr(gnd,prd)[0];
        print('\t \t Fixed point correlation: %.3f' %p_cor)
    else:
        p_cor = np.nan;
        
    for idx in range(num_iters):
        theta_old = theta.copy()
        # Compute predictions and prediction errors
        dhat      = np.tanh(x @ theta);
        if solver_type == 'weighted':
            E_pred    = (d-dhat) /(1-d**2); 
            scale = np.sum(abs(E_pred)>threshold,0);
            
            if verbose:
                per_not_used = np.mean(scale) / T_data;
                
            scale = T_data/(T_data-scale);
            
            E_pred[abs(E_pred)>threshold] = 0
            E_pred = E_pred * scale;
        elif solver_type == 'standard':
            E_pred    = (d-dhat);
            per_not_used = 0;
        elif solver_type == 'robust':
            E_pred    = (d-dhat) /(1-d**2);
            per_not_used = 0;
            E_pred[abs(E_pred)>threshold] = threshold * np.sign(E_pred[abs(E_pred)>threshold])

        
        
        # Perform the first primal variable update
        theta     = Xm @ theta + Xp @ E_pred;
        conv = np.sqrt(n_rec) * np.sqrt(np.mean((theta - theta_old ) **2 ))
       
        
        if ( (verbose == 2) & (gnd is not None) ) :
            prd = (theta[:n_rec,:].T).flatten();
            p_cor = pearsonr(gnd,prd)[0];
        else:
            p_cor = np.nan;
        
            
            
        if ((verbose == 2) & (np.mod(idx,10) == 9)):
            temp = time_now.localtime()
            current_time = time_now.strftime("%H:%M:%S", temp)
            slope = (gnd @ prd) / (gnd@gnd) 
            rmse  = np.sqrt(np.mean( (prd-gnd)**2 ))
            print('\t \t %s: Iteration %d finished. RMSE: %.4f. Correlation %.3f. Slope %.3f. Not used %.2f. Convergence %.7f.' \
                  %(current_time,idx + 1,rmse,p_cor,slope,100*per_not_used,conv))
                
        if ((check_convergence>0) & (idx > 10)):
            if conv < 1e-5:
                break
    
    if verbose:
        temp = time_now.localtime()
        current_time = time_now.strftime("%H:%M:%S", temp)
        tt = time_now.perf_counter() - start_time
        print('%s: CoRNN solver finished in %.2f mins' %(current_time,tt/60))

    return theta
    


def solve_gradient_descent(r_in,r_out,u_in,alph = 0.1,
                         l2 = 1e-5,verbose = 0,
                         initialize_fp = 0,num_iters = 1e4,
                         threshold = 1,learning_rate = 0.001,
                         gnd = None,momentum = 0,solver_type = 'weighted'):
    
    if verbose:
        temp = time_now.localtime()
        current_time = time_now.strftime("%H:%M:%S", temp)
        print('%s: Running GD solver.' %(current_time))
    start_time = time_now.perf_counter()
    
    T_data = r_in.shape[0]
    n_rec = r_in.shape[1]
    if u_in is not None:
        n_in = u_in.shape[1]
    else:
        n_in = 0
    n_tot = n_rec + n_in;
    
    # Get the inputs x and targets d
    if n_in >0:
        x = np.c_[r_in,u_in];
    else:
        x = r_in.copy()
    d = (r_out - (1-alph)*r_in)/alph
    d[d<=-1+1e-6] = -1 + 1e-6;
    d[d>=1-1e-6]  = 1-1e-6;
    

    # Scale the regularization and compute fixed point
    l2_scaled  = l2*T_data;
    if initialize_fp:
        A = x.T @ x + (l2_scaled) * np.diag(np.ones(x.shape[1]));
        
        
        Ainv = inv(A);
        Xp   = Ainv @ x.T;
        z    = np.arctanh(d);
        theta_fp = Xp @ z;
        
    if initialize_fp:
        theta = theta_fp
    else:
        theta = np.zeros([n_tot,n_rec ])
    
    for idx in range(num_iters):
            
        
        grad = compute_gradient(x,d,theta,l2,threshold,solver_type)
        del_gd = - grad * learning_rate ;
        if idx == 0:
            del_gd_old = del_gd.copy()
            
        delta_gd = momentum * (del_gd_old) + (1- momentum) * del_gd
        
        del_gd_old = delta_gd.copy()
        
        theta = theta + delta_gd
        if ((verbose == 2) & (np.mod(idx,300) == 299) ):
            prd = (theta[:n_rec,:].T).flatten();
            if gnd is not None:
                p_cor = pearsonr(gnd,prd)[0];
            else:
                p_cor = np.nan
            temp = time_now.localtime()
            current_time = time_now.strftime("%H:%M:%S", temp)
            rmse  = np.sqrt(np.mean( (prd-gnd)**2 ))
            slope = (gnd @ prd) / (gnd@gnd)
            print('\t %s: Iteration %d finished. RMSE: %.4f. Correlation %.3f. Slope %.3f.' \
                  %(current_time,idx + 1,rmse,p_cor,slope))
   
    
    if verbose:
        temp = time_now.localtime()
        current_time = time_now.strftime("%H:%M:%S", temp)
        tt = time_now.perf_counter() - start_time
        print('%s: gradient descent finished in %.2f mins' %(current_time,tt/60))
    
    return theta   
        
        

def solve_newton_descent(r_in,r_out,u_in,alph = 0.1,
                         l2 = 1e-5,verbose = 0,
                         initialize_fp = 0,num_iters = 10,
                         threshold = 1,solver_type = 'weighted'):
    if verbose:
        temp = time_now.localtime()
        current_time = time_now.strftime("%H:%M:%S", temp)
        print('%s: Running NT solver.' %(current_time))
    start_time = time_now.perf_counter()
    
    T_data = r_in.shape[0]
    n_rec = r_in.shape[1]
    if u_in is not None:
        n_in = u_in.shape[1]
    else:
        n_in = 0
    n_tot = n_rec + n_in;
    theta = np.zeros([n_tot,n_rec]);
    
    # Get the inputs x and targets d
    if n_in >0:
        x = np.c_[r_in,u_in];
    else:
        x = r_in.copy()
    d = (r_out - (1-alph)*r_in)/alph
    d[d<=-1+1e-6] = -1 + 1e-6;
    d[d>=1-1e-6]  = 1-1e-6;
    

    # Scale the regularization and compute fixed point
    l2_scaled  = l2*T_data;
    if initialize_fp:
        A = x.T @ x + (l2_scaled) * np.diag(np.ones(x.shape[1]));
        
        
        Ainv = inv(A);
        Xp   = Ainv @ x.T;
        z    = np.arctanh(d);
        theta_fp = Xp @ z;
    
    num_cores = np.array(multiprocessing.cpu_count()/2).astype(int)
    if verbose:
        temp = time_now.localtime()
        current_time = time_now.strftime("%H:%M:%S", temp)
        print('%s: NT-descent runs with %d cores.' %(current_time,num_cores))
    
    def solve_descent(iters):
        if initialize_fp:
            theta_in = theta_fp[:,iters].copy()
        else:
            theta_in = None
        beta = run_nt_algorithm(x,d[:,iters],num_iters,l2,theta_in,threshold,solver_type)
        return beta
    
    (result) = Parallel(n_jobs=num_cores)(delayed(solve_descent)(iters) for iters in range(n_rec))
    
    theta = np.array(result).T
    
    if verbose:
        temp = time_now.localtime()
        current_time = time_now.strftime("%H:%M:%S", temp)
        tt = time_now.perf_counter() - start_time
        print('%s: NT-descent finished in %.2f mins' %(current_time,tt/60))
    
    return theta
    
def run_nt_algorithm(x,d_tar,n_iter,l2,theta_in,threshold,solver_type):
    
    if theta_in is not None:
        beta = theta_in
    else:
        beta = np.zeros([x.shape[1]])

    for idx in range(n_iter):
       
        
        grad = compute_gradient(x,d_tar,beta,l2,threshold,solver_type)
        
        hes  = compute_hessian(x,d_tar,beta,l2,threshold,solver_type)
        del_nt = -solve(hes, grad,assume_a='pos')
        kappa = 1
        gamma = 0.4;
        chi = 0.6;
        count = 0
        
        while(   compute_loss_function(x,d_tar,beta + kappa * del_nt,l2) >=  \
              compute_loss_function(x,d_tar,beta,l2) + gamma * kappa* (grad @ del_nt)  ):
    
            kappa = chi * kappa
            count = count + 1
            if count == 4:
                kappa = 1;
                break
            
        delta_gd = kappa * del_nt
        
        beta = beta + delta_gd
        
        
            
    return beta

def compute_gradient(x,d_tar,beta,reg,threshold,solver_type):
    T_data = x.shape[0]
    
    
    
    dhat = np.tanh( x @ beta )
    
    if solver_type == 'weighted':
        E_pred =  ( (dhat-d_tar) / (1-d_tar**2)  ).T;
        scale = np.sum(abs(E_pred)>threshold,0);
        if np.max(scale) > 0.9 * T_data:
            scale = np.sum(abs(E_pred)>10*threshold,0);
            threshold = 10* threshold;
        scale = T_data/(T_data-scale);
        
        E_pred[abs(E_pred)>threshold] = 0
        E_pred = E_pred * scale;
        E_pred = E_pred @ x / T_data
    else:
        E_pred =  ( (dhat-d_tar) ).T;
        E_pred = E_pred @ x / T_data;
    
    gradient = E_pred.T  + reg * beta
    
    
    return gradient

def compute_hessian(x,d_tar,beta,reg,threshold,solver_type):
    T_data = x.shape[0]
    
    dhat = np.tanh( x @ beta )
    
    if solver_type == 'weighted':
        E_pred =  ( (dhat-d_tar) / (1-d_tar**2)  );
        scale = np.sum(abs(E_pred)>threshold,0);
        if np.max(scale) > 0.9 * T_data:
            scale = np.sum(abs(E_pred)>10*threshold,0);
            threshold = 10* threshold;
        scale = T_data/(T_data-scale);
        
        temp = np.sqrt(1-dhat**2) / np.sqrt(1-d_tar**2)
        temp[abs(E_pred)>threshold] = 0
        temp = temp * scale;
    else:
        temp = np.sqrt(1-dhat**2);
    
    r = x * temp[:,None];
    
    
    H   = r.T @ r /r.shape[0] + reg * np.diag(np.ones(r.shape[1]))    
    
    return H
        
        

def compute_loss_function(x,d,beta,reg):
    
    dhat = np.tanh( x @ beta )
    dhat[dhat>=1] = 1-1e-16
    dhat[dhat<=-1] = -1+1e-16
    loss = - np.mean( 0.5*(d+1)* np.log(0.5*(dhat+1)) +  0.5*(1-d)* np.log(0.5*(1-dhat))   ) + reg * np.sum(beta**2)/2

    return loss


def solve_pytorch(r_in,r_out,u_in,alph = 0.1,
                         l2 = 1e-5,verbose = 0,
                         initialize_fp = 0,num_iters = 1e4,
                         threshold = 2,learning_rate = 0.001,
                         gnd = None,solver_type = 'logistic'):
    if verbose:
        temp = time_now.localtime()
        current_time = time_now.strftime("%H:%M:%S", temp)
        print('%s: Running Pytorch-GD solver.' %(current_time))
    start_time = time_now.perf_counter()
    
    n_rec = r_in.shape[1]
    if u_in is not None:
        n_in = u_in.shape[1]
    else:
        n_in = 0
    
    # Get the inputs x and targets d
    if n_in >0:
        x = np.c_[r_in,u_in];
    else:
        x = r_in.copy()
    d = (r_out - (1-alph)*r_in)/alph
    d[d<=-1+1e-6] = -1 + 1e-6;
    d[d>=1-1e-6]  = 1-1e-6;
    if initialize_fp:
        T_data = x.shape[0]
        A = x.T @ x + l2 *T_data *1e3* np.diag(np.ones(x.shape[1]));
        
        
        Ainv = inv(A);
        Xp   = Ainv @ x.T;
        z    = np.arctanh(d);
        theta_fp = Xp @ z;

    
    # Define the network
    class Model(nn.Module):
        def __init__(self, n_rec, n_in):
            super(Model, self).__init__()
            self.n_rec = n_rec
            self.n_in = n_in
            self.linear = nn.Linear(n_rec + n_in, n_rec,bias=False)
            self.tanh = nn.Tanh()
        
        def forward(self, x_in):
            #x_in = torch.tensor(x_in, dtype=torch.float32)
            d_out = self.tanh(self.linear(x_in))
            return d_out
        
    d = torch.tensor(d, dtype=torch.float32)
    x = torch.tensor(x, dtype=torch.float32)
    model = Model(n_rec, n_in)
    # define the multi target logistic regression loss function
    if solver_type == 'logistic':
        criterion = nn.BCELoss()
    else:
        criterion = nn.MSELoss()
    optimizer = optim.Adam(model.parameters(), lr=learning_rate, weight_decay=l2)
    if initialize_fp:
        model.linear.weight.data = torch.tensor(theta_fp.T, dtype=torch.float32)
    for idx in range(num_iters):
        d_out = model(x)

        # compute the loss
        loss = criterion((1 + d_out)/2, (1 + d)/2)
        
        # backward
        loss.backward()
        
        
        # update the weights
        optimizer.step()
        # zero the gradients
        optimizer.zero_grad()
        

        if ((verbose == 2) & (np.mod(idx,300) == 299) ):
            weights = model.linear.weight
            prd = weights[:, :n_rec].detach().numpy().flatten()
            if gnd is not None:
                p_cor = pearsonr(gnd,prd)[0];
            else:
                p_cor = np.nan
            slope = (gnd @ prd) / (gnd@gnd)
            
            temp = time_now.localtime()
            current_time = time_now.strftime("%H:%M:%S", temp)
            rmse  = np.sqrt(np.mean( (prd-gnd)**2 ))
            print('\t %s: Iteration %d finished. RMSE: %.4f Correlation %.3f. Slope %.3f.' \
                  %(current_time,idx + 1,rmse,p_cor,slope))
   
    
    if verbose:
        temp = time_now.localtime()
        current_time = time_now.strftime("%H:%M:%S", temp)
        tt = time_now.perf_counter() - start_time
        print('%s: Pytorch finished in %.2f mins' %(current_time,tt/60))
    
    return model.linear.weight.detach().numpy().T


def solve_pytorch_gpu(r_in,r_out,u_in,alph = 0.1,
                         l2 = 1e-5,verbose = 0,
                         initialize_fp = 0,num_iters = 1e4,
                         threshold = 2,learning_rate = 0.001,
                         gnd = None,solver_type = 'logistic'):
    if verbose:
        temp = time_now.localtime()
        current_time = time_now.strftime("%H:%M:%S", temp)
        print('%s: Running Pytorch-GD solver.' %(current_time))
    start_time = time_now.perf_counter()
    
    n_rec = r_in.shape[1]
    if u_in is not None:
        n_in = u_in.shape[1]
    else:
        n_in = 0

    try:
        gpu_device = torch.device("cuda:0");
        a = torch.tensor(1,device = gpu_device);
    except:
        gpu_device = torch.device("mps");
    
    # Get the inputs x and targets d
    if n_in >0:
        x = np.c_[r_in,u_in];
    else:
        x = r_in.copy()
    d = (r_out - (1-alph)*r_in)/alph
    d[d<=-1+1e-6] = -1 + 1e-6;
    d[d>=1-1e-6]  = 1-1e-6;

    # Define the network
    class Model(nn.Module):
        def __init__(self, n_rec, n_in):
            super(Model, self).__init__()
            self.n_rec = n_rec
            self.n_in = n_in
            self.linear = nn.Linear(n_rec + n_in, n_rec,bias=False)
            self.tanh = nn.Tanh()
        
        def forward(self, x_in):
            #x_in = torch.tensor(x_in, dtype=torch.float32)
            d_out = self.tanh(self.linear(x_in))
            return d_out
    


    d = torch.tensor(d, dtype=torch.float32,device = gpu_device)
    x = torch.tensor(x, dtype=torch.float32,device = gpu_device)

    if initialize_fp:
        T_data = x.shape[0]
        n_tot = n_rec+ n_in
        reg_term = torch.tensor((l2) * T_data * 1e3 * np.diag(np.ones(n_tot)),\
                                device=gpu_device,dtype = torch.float32)

        A = x.T @ x + reg_term;
        
        try:
            Ainv = torch.linalg.inv(A);
        except:
            Ainv = torch.linalg.inv(A.cpu()).to(gpu_device);
        Xp   = Ainv @ x.T;
        z    = torch.arctanh(d);
        theta_fp = Xp @ z;
        theta_fp = theta_fp.cpu().detach().numpy()
        
    model = Model(n_rec, n_in)
    model.to(gpu_device)
    # define the multi target logistic regression loss function
    if solver_type == 'logistic':
        criterion = nn.BCELoss()
    else:
        criterion = nn.MSELoss()
    optimizer = optim.Adam(model.parameters(), lr=learning_rate, weight_decay=l2)
    if initialize_fp:
        model.linear.weight.data = torch.tensor(theta_fp.T, dtype=torch.float32).to(gpu_device)
    for idx in range(num_iters):
        d_out = model(x)

        # compute the loss
        loss = criterion((1 + d_out)/2, (1 + d)/2)
        
        # backward
        loss.backward()
        
        
        # update the weights
        optimizer.step()
         
        # zero the gradients
        optimizer.zero_grad()
        

        if ((verbose == 2) & (np.mod(idx,300) == 299)):
            weights = model.linear.weight
            prd = weights[:, :n_rec].cpu().detach().numpy().flatten()
            if gnd is not None:
                p_cor = pearsonr(gnd,prd)[0];
            else:
                p_cor = np.nan
            slope = (gnd @ prd) / (gnd@gnd)
            
            temp = time_now.localtime()
            current_time = time_now.strftime("%H:%M:%S", temp)
            rmse  = np.sqrt(np.mean( (prd-gnd)**2 ))
            print('\t %s: Iteration %d finished. RMSE: %.4f Correlation %.3f. Slope %.3f.' \
                  %(current_time,idx + 1,rmse,p_cor,slope))
   
    
    if verbose:
        temp = time_now.localtime()
        current_time = time_now.strftime("%H:%M:%S", temp)
        tt = time_now.perf_counter() - start_time
        print('%s: Pytorch finished in %.2f mins' %(current_time,tt/60))
    
    return model.linear.weight.cpu().detach().numpy().T
 

def fit_FORCE(r,u=None,alph = 0.1,
                lam = 100,lam_in = 1,g_in = 3,verbose = 0,
                initialize_fp = 0,num_iters = 1e3,
                gnd = None,solver_type = 'currents'):
    if verbose:
        temp = time_now.localtime()
        current_time = time_now.strftime("%H:%M:%S", temp)
        print('%s: Running FORCE solver on %s.' %(current_time,solver_type))
    
    def predict_single(r_before,u_in = None):
        if u_in is not None:
            x = np.dot(w_rec,r_before) + np.dot(w_in,u_in);
        else:
            x = np.dot(w_rec,r_before);
        rout = (1-alph) * r_before + alph * np.tanh(x)
        
        return rout,x
    
    if initialize_fp:
        if u is not None:
            n_in = u.shape[1]
        else:
            n_in = 0
        r_in = r[:-1,:]
        r_out = r[1:,:]
        d = (r_out - (1-alph)*r_in)/alph
        d[d<=-1+1e-6] = -1 + 1e-6;
        d[d>=1-1e-6]  = 1-1e-6;
        if n_in >0:
            x = np.c_[r_in,u];
        else:
            x = r_in.copy()
        T_data = x.shape[0]
        A = x.T @ x + lam/(10*T_data) * np.diag(np.ones(x.shape[1]));
        
        
        Ainv = inv(A);
        Xp   = Ainv @ x.T;
        z    = np.arctanh(d);
        theta_fp = Xp @ z;

    
    T = r.shape[0]-1
    n_rec=  r.shape[1]
    if u is not None and u.shape[0] != T:
        raise Exception('There is a missmatch between u and r dimensions!!!')
    
   
    
    temp=np.random.normal(0,g_in/np.sqrt(n_rec),n_rec*n_rec).reshape(n_rec,n_rec)
    np.fill_diagonal(temp,0)
    w_rec = temp
    
    if u is not None:
        n_in = u.shape[1]
        w_in =np.random.normal(0,g_in/np.sqrt(n_in),n_in*n_rec).reshape(n_rec,n_in)
    else:
        n_in = 0
    
    if initialize_fp:
        w_rec = theta_fp[:n_rec,:].T
        if n_in >0:
            w_in = theta_fp[n_rec:,:].T.reshape(n_rec,n_in)
        if (verbose == 2):
            prd = w_rec.flatten()
            if gnd is not None:
                p_cor = pearsonr(gnd,prd)[0];
            else:
                p_cor = np.nan
            print('\t %s: FP initialized: Correlation %.3f.' \
                  %(current_time,p_cor))
    
    
    for k in range(num_iters):
        P = np.diag(np.ones(n_rec))/lam;
        if n_in > 0:
            P_in = np.diag(np.ones(n_in))/lam_in;
        r_bef = r[0,:];
        for i in range(T):
            temp = (r[i+1,:] - (1-alph)*r[i,:])/alph
            temp[temp>=1-1e-6] = 1-1e-6
            temp[temp<=-1+1e-6] = -1+1e-6
            
            x_now = np.arctanh(temp)
            if n_in > 0:
                r_out,x_out = predict_single(r_bef,u[i])
            else:
                r_out,x_out = predict_single(r_bef)
            
            if solver_type == 'currents':
                e_min = x_out - x_now;
            elif solver_type == 'firing_rates':
                e_min = (r_out - r[i+1,:])/alph;
                
            Pxr   = P @ r_bef;
            rxPxr = r_bef @ (P @ r_bef) + 1
            delP  = - np.outer(Pxr, Pxr) / rxPxr
            P     = P + delP;
            delW  = - np.outer(e_min,  P @ r_bef)
            
            
            w_rec = w_rec + delW;
            
            if n_in > 0:
                Pxu = P_in @ u[i]
                uxPxu = u[i] @ (P_in @ u[i]) + 1
                delP  = - np.outer(Pxu, Pxu) / uxPxu
                P_in     = P_in + delP;
                delW  = - np.outer(e_min,  P_in @ u[i])
                w_in = w_in + delW;
                r_bef,x_pred = predict_single(r_bef,u[i])
            else:
                r_bef,x_pred = predict_single(r_bef)
            
            if solver_type == 'currents':
                e_pls = x_pred - x_now
            elif solver_type == 'firing_rates':
                e_pls = (r_bef - r[i+1,:])/alph;
            
        if ((verbose == 2)):
            prd = w_rec.flatten()
            if gnd is not None:
                p_cor = pearsonr(gnd,prd)[0];
            else:
                p_cor = np.nan
            slope = (gnd @ prd) / (gnd@gnd)
            
            temp = time_now.localtime()
            current_time = time_now.strftime("%H:%M:%S", temp)
            rmse  = np.sqrt(np.mean( (prd-gnd)**2 ))
            print('\t %s: Iteration %d finished. RMSE: %.4f Correlation %.3f. Slope %.3f. Conv. %.3f.' \
                  %(current_time,k + 1,rmse,p_cor,slope,np.min(e_pls/e_min)))
    
    if n_in>0:
        theta = np.c_[w_rec,w_in]
    else:
        theta = w_rec
    
    return theta.T

 
def solve_corrn_gpu(r_in,r_out,u_in = None, alph = 0.1,  float_type = '32bit',
                l2 = 1e-4, threshold = 0.2,initialize_fp = 1,
                verbose = 0,mask = None, check_convergence = 0,
                num_iters = 30,gnd = None,solver_type = 'weighted'):
    if verbose:
        temp = time_now.localtime()
        current_time = time_now.strftime("%H:%M:%S", temp)
        print('%s: Running CoRNN solver.' %(current_time))
    start_time = time_now.perf_counter()
    try:
        gpu_device = torch.device("cuda:0");
        a = torch.tensor(1,device = gpu_device);
    except:
        gpu_device = torch.device("mps");
    num_iters = int(num_iters);
    T_data = r_in.shape[0]
    n_rec = r_in.shape[1]
    if u_in is not None:
        n_in = u_in.shape[1]
    else:
        n_in = 0
    
    # Get the inputs x and targets d
    if n_in >0:
        x = np.c_[r_in,u_in];
    else:
        x = r_in.copy()
    d = (r_out - (1-alph)*r_in)/alph
    d[d<=-1+1e-6] = -1 + 1e-6;
    d[d>=1-1e-6]  = 1-1e-6;
    if float_type == '32bit':
        dtype = torch.float32
    else:
        dtype = torch.float64
    n_tot = x.shape[1]
    
    x = torch.tensor(x,device=gpu_device, dtype=dtype)
    d = torch.tensor(d,device=gpu_device, dtype=dtype)
    
    if verbose == 2:
        temp = time_now.localtime()
        current_time = time_now.strftime("%H:%M:%S", temp)
        print('\t %s: Performing initial inverse operations' %(current_time))
    
    # Scale the regularization and compute the fixed point
    l2  = l2*T_data;
    
    
    
    if mask is None:
        mask = np.r_[np.diag(np.ones(n_rec)),np.zeros([n_in,n_rec]) ]
        mask = (mask == 1)
        
    reg_term = torch.tensor((l2) * np.diag(np.ones(n_tot)),\
                            device=gpu_device, dtype=dtype)

    A = x.T @ x + reg_term;
    
    
    try:
        Ainv = torch.linalg.inv(A);
    except:
        Ainv = torch.linalg.inv(A.cpu()).to(gpu_device);
    Xp   = Ainv @ x.T;
    Xm  = Xp @ x; 
    z    = torch.arctanh(d);
    theta_fp = Xp @ z;
    if initialize_fp:
        theta = theta_fp;
    else:
        w_rec = np.random.normal(0,1/np.sqrt(n_rec),n_rec*n_tot).reshape(n_tot,n_rec)
        theta = torch.tensor(w_rec,device=gpu_device, dtype=dtype);
    if ( (verbose == 2) & (gnd is not None) ) :
        prd = (theta[:n_rec,:].T).flatten().cpu().numpy();
        p_cor = pearsonr(gnd,prd)[0];
        print('\t \t Fixed point correlation: %.3f' %p_cor)
    else:
        p_cor = np.nan;
    
    if verbose == 2:
        temp = time_now.localtime()
        current_time = time_now.strftime("%H:%M:%S", temp)
        print('\t %s: Starting iterations' %(current_time))
        
    for idx in range(num_iters):
        # Compute predictions and prediction errors
        dhat      = torch.tanh(x @ theta);
        if solver_type == 'weighted':
            E_pred    = (d-dhat) /(1-d**2); 
            per_not_used = torch.sum(abs(E_pred)>threshold) / (n_rec * T_data)
            E_pred[abs(E_pred)>threshold] = 0       
        elif solver_type == 'standard':
            E_pred    = (d-dhat);
            per_not_used = 0;
        elif solver_type == 'robust':
            E_pred    = (d-dhat) /(1-d**2);
            per_not_used = 0;
            E_pred[abs(E_pred)>threshold] = threshold * np.sign(E_pred[abs(E_pred)>threshold])
            
            
       
        
        
        
        
        # Perform the first primal variable update
        theta     = Xm @ theta + Xp @ E_pred
       
        
        if ( (verbose == 2) & (gnd is not None) ) :
            prd = (theta[:n_rec,:].T).flatten().cpu().numpy();
            p_cor = pearsonr(gnd,prd)[0];
        else:
            p_cor = np.nan;
        
            
            
        if ((verbose == 2) & (np.mod(idx,300) == 299) ):
            temp = time_now.localtime()
            current_time = time_now.strftime("%H:%M:%S", temp)
            slope = (gnd @ prd) / (gnd@gnd) 
            rmse  = np.sqrt(np.mean( (prd-gnd)**2 ))
            print('\t \t %s: Iteration %d finished. RMSE: %.4f. Correlation %.3f. Slope %.3f. Not used %.2f.' \
                  %(current_time,idx + 1,rmse,p_cor,slope,100*per_not_used))
                
    
    if verbose:
        temp = time_now.localtime()
        current_time = time_now.strftime("%H:%M:%S", temp)
        tt = time_now.perf_counter() - start_time
        print('%s: CoRNN solver finished in %.2f mins' %(current_time,tt/60))

    return theta.cpu().numpy()

def fit_FORCE_gpu(r,u=None,alph = 0.1,
                lam = 100,lam_in = 1,g_in = 3,verbose = 0,
                initialize_fp = 0,num_iters = 1e3,
                gnd = None,solver_type = 'currents'):
    if verbose:
        temp = time_now.localtime()
        current_time = time_now.strftime("%H:%M:%S", temp)
        print('%s: Running FORCE solver on %s.' %(current_time,solver_type))
    
    try:
        gpu_device = torch.device("cuda:0");
        a = torch.tensor(1,device = gpu_device);
    except:
        gpu_device = torch.device("mps");
    
    def predict_single(r_before,u_in = None):
        if u_in is not None:
            x = w_rec @ r_before +w_in @u_in;
        else:
            x = w_rec @r_before;
        rout = (1-alph) * r_before + alph * torch.tanh(x)
        
        return rout,x
    
    if initialize_fp:
        if u is not None:
            n_in = u.shape[1]
        else:
            n_in = 0
        r_in = r[:-1,:]
        r_out = r[1:,:]
        d = (r_out - (1-alph)*r_in)/alph
        d[d<=-1+1e-6] = -1 + 1e-6;
        d[d>=1-1e-6]  = 1-1e-6;
        if n_in >0:
            x = np.c_[r_in,u];
        else:
            x = r_in.copy()
            
        x = torch.tensor(x,device=gpu_device,dtype = torch.float32)
        d = torch.tensor(d,device=gpu_device,dtype = torch.float32)
        T_data = x.shape[0]
       
        reg_term = torch.tensor(lam/T_data * np.diag(np.ones(x.shape[1])),\
                                device=gpu_device,dtype = torch.float32)

        A = x.T @ x + reg_term;
        
        
        try:
            Ainv = torch.linalg.inv(A);
        except:
            Ainv = torch.linalg.inv(A.cpu()).to(gpu_device);
        Xp   = Ainv @ x.T;
        z    = torch.arctanh(d);
        theta_fp = Xp @ z;
    
    
    r = torch.tensor(r,device=gpu_device,dtype = torch.float32)
    alph = torch.tensor(alph,device=gpu_device,dtype = torch.float32)
    T = r.shape[0]-1
    n_rec=  r.shape[1]
    if u is not None and u.shape[0] != T:
        raise Exception('There is a missmatch between u and r dimensions!!!')
    
   
    
    temp=np.random.normal(0,g_in/np.sqrt(n_rec),n_rec*n_rec).reshape(n_rec,n_rec)
    np.fill_diagonal(temp,0)
    w_rec = torch.tensor(temp,device=gpu_device,dtype = torch.float32)
    
    if u is not None:
        n_in = u.shape[1]
        w_in =np.random.normal(0,g_in/np.sqrt(n_in),n_in*n_rec).reshape(n_rec,n_in)
        w_in = torch.tensor(w_in,device=gpu_device,dtype = torch.float32)
    else:
        n_in = 0
    
    if initialize_fp:
        w_rec = theta_fp[:n_rec,:].clone().detach().T
        if n_in >0:
            w_in = torch.tensor(theta_fp[n_rec:,:].clone().detach().T.reshape(n_rec,n_in),device=gpu_device,dtype = torch.float32)
        if (verbose == 2):
            prd = w_rec.flatten().cpu().numpy();
            if gnd is not None:
                p_cor = pearsonr(gnd,prd)[0];
            else:
                p_cor = np.nan
            print('\t %s: FP initialized: Correlation %.3f.' \
                  %(current_time,p_cor))
    
    
    for k in range(num_iters):
        P = torch.tensor(np.diag(np.ones(n_rec))/lam,device=gpu_device,dtype = torch.float32);
        if n_in > 0:
            P_in = torch.tensor(np.diag(np.ones(n_in))/lam_in,device=gpu_device,dtype = torch.float32);
        r_bef = r[0,:];
        for i in range(T):
            temp = (r[i+1,:] - (1-alph)*r[i,:])/alph
            temp[temp>=1-1e-6] = 1-1e-6
            temp[temp<=-1+1e-6] = -1+1e-6
            
            x_now = torch.arctanh(temp)
            if n_in > 0:
                r_out,x_out = predict_single(r_bef,u[i])
            else:
                r_out,x_out = predict_single(r_bef)
            
            if solver_type == 'currents':
                e_min = x_out - x_now;
            elif solver_type == 'firing_rates':
                e_min = (r_out - r[i+1,:])/alph;
                
            Pxr   = P @ r_bef;
            rxPxr = r_bef @ (P @ r_bef) + 1
            delP  = - torch.outer(Pxr, Pxr) / rxPxr
            P     = P + delP;
            delW  = - torch.outer(e_min,  P @ r_bef)
            
            
            w_rec = w_rec + delW;
            
            if n_in > 0:
                Pxu = P_in @ u[i]
                uxPxu = u[i] @ (P_in @ u[i]) + 1
                delP  = - torch.outer(Pxu, Pxu) / uxPxu
                P_in     = P_in + delP;
                delW  = - torch.outer(e_min,  P_in @ u[i])
                w_in = w_in + delW;
                r_bef,x_pred = predict_single(r_bef,u[i])
            else:
                r_bef,x_pred = predict_single(r_bef)
            
            if solver_type == 'currents':
                e_pls = x_pred - x_now
            elif solver_type == 'firing_rates':
                e_pls = (r_bef - r[i+1,:])/alph;
            
        if ((verbose == 2)):
            prd =  w_rec.flatten().cpu().numpy();
            if gnd is not None:
                p_cor = pearsonr(gnd,prd)[0];
            else:
                p_cor = np.nan
            slope = (gnd @ prd) / (gnd@gnd)
            
            temp = time_now.localtime()
            current_time = time_now.strftime("%H:%M:%S", temp)
            rmse  = np.sqrt(np.mean( (prd-gnd)**2 ))
            print('\t %s: Iteration %d finished. RMSE: %.4f Correlation %.3f. Slope %.3f. Conv. %.3f.' \
                  %(current_time,k + 1,rmse,p_cor,slope,torch.min(e_pls/e_min)))
    
    w_rec = w_rec.cpu().numpy()
    
    if n_in>0:
        w_in = w_in.cpu().numpy()
        theta = np.c_[w_rec,w_in]
    else:
        theta = w_rec
    
    return theta.T

 
```
