## Supplementary material for "Partial observation can induce mechanistic mismatches in data-constrained models of neural dynamics": Code: utils_RNN.py

```
import numpy as np

from simulation_conditions import *
def get_stationary_cov(B, c = 0.0):
    D = len(B)
    I = np.eye(D)
    J = B - I
    noise_cov = (1-c)*I+c*np.ones((D,D))
    cov = scipy.linalg.solve_continuous_lyapunov(J, -noise_cov)
    return cov

def get_stationary_fit(B, N_obs, cov, ff_chain = False):
    if ff_chain: # use for higher precision, as only the last row is nonzero
        learned = generate_jordan_block(N_obs)
        final_row = np.linalg.solve(cov[:N_obs, :N_obs], (B @ cov)[N_obs - 1, :N_obs])
        learned[-1] = final_row
    else:
        learned = (B @ cov)[:N_obs, :N_obs] @ np.linalg.inv(cov[:N_obs, :N_obs])
    return learned

def get_learned_stationary(B, N_obs):
    cov = get_stationary_cov(B)
    return get_stationary_fit(B, N_obs, cov)

def overlap(x, y):
    L = min(len(x), len(y))
    x_L = x[:L].copy()
    x_L /= np.sqrt(np.vdot(x_L, x_L))
    y_L = y[:L].copy()
    y_L /= np.sqrt(np.vdot(y_L, y_L))
    return np.abs(np.real(np.vdot(x_L, y_L)))

def get_flow_field(w, pca, x_range = [-75, 75], y_range= None, x_num = 10, pc_pad = None):
    dx = (x_range[1] - x_range[0]) / x_num
    if y_range is None:
        y_range = x_range
    dy = (y_range[1] - y_range[0]) / x_num
    x,y = np.meshgrid(np.linspace(x_range[0], x_range[1]+dx, x_num),np.linspace(y_range[0], y_range[1]+dy, x_num))
    xy = np.vstack([x.ravel(), y.ravel()]).T
    if pc_pad is None:
        pc_pad = np.zeros((len(xy),len(w) - 2))
    else:
        pc_pad = np.tile(pc_pad, (len(xy), 1))
    xy = np.hstack([xy,  pc_pad])
    xy_orig = pca.inverse_transform(xy) #(x_num, N_obs)
    dim = len(w)
    I = np.eye(dim)
    flow = (w-I) @ xy_orig.T #(N_obs, x_num)
    flow_pca = pca.transform(flow.T)
    return xy, flow_pca[:, :2]

def henrici_nonnormality(A):
    evals = np.linalg.eigvals(A)
    norm = np.linalg.norm(A)
    return np.sqrt(np.abs(norm **2 - np.sum(np.abs(evals)**2)))/norm

def get_subsampling_curves(n_neuron, interval = 1, top_n=5,ff_chain = False, detailed = True, **kwargs):
    B, _ = get_W(n_neuron, **kwargs)
    h_vals = []
    evals = []
    svals = []
    uvecs_overlap = []
    vvecs_overlap = []
    evecs_overlap = []
    U_orig, s, Vh_orig = np.linalg.svd(B)
    evs, evecs_orig, tau = get_eigenproperties(B)
    las = []
    cov = get_stationary_cov(B)
    for i in range(1, n_neuron + 1, interval):
        A_hat = get_stationary_fit(B, i, cov, ff_chain=ff_chain)
        h_vals.append(henrici_nonnormality(A_hat))
        evs, evecs, tau = get_eigenproperties(A_hat)
        if detailed:
            U_hat, s_hat, Vh_hat = np.linalg.svd(A_hat)
            svals.append(np.concatenate([s_hat[:top_n], np.zeros(top_n - len(s_hat[:top_n]))]))
            uvecs_overlap.append(overlap(U_orig[:, 0], U_hat[:, 0]))
            vvecs_overlap.append(overlap(Vh_hat[0], Vh_orig[0]))
            evecs_overlap.append(overlap(evecs[:, 0], evecs_orig[:, 0]))

        evals.append(np.concatenate([np.real(evs[:top_n]), np.zeros(top_n - len(evs[:top_n]))]))
        if i > 1:
            las.append(np.log2(tau[0] / tau[1]))
        else:
            las.append(0)
    return B, np.array(h_vals), np.array(evals), np.array(las), np.array(
        svals), np.array(uvecs_overlap), np.array(vvecs_overlap), np.array(evecs_overlap)

def participation_ratio(cov):
    evals = np.linalg.eigvals(cov)
    return np.sum(evals) ** 2 / np.sum(evals**2)

def explained_var(X_gt, X_hat, phi= lambda x: x):
    num = np.mean((phi(X_hat) - phi(X_gt))**2)
    mean_gt = np.mean(phi(X_gt), axis=0, keepdims=True)
    denom = np.mean((phi(X_gt) - mean_gt)**2)
    return 1 - num/denom
```
